# Disentangling structure-function relationships between the human hippocampus and the whole brain using track-weighted dynamic functional connectivity

**DOI:** 10.1101/2025.10.19.683338

**Authors:** Marshall A. Dalton, Jinglei Lv, Yifei Sun, Arkiev D’Souza, Fernando Calamante

## Abstract

Understanding how structural and functional connectivity shape hippocampal interactions with the rest of the brain is critical for elucidating its role in cognition. Here, we combine high-resolution diffusion MRI, a novel fibre-tracking pipeline designed to specifically probe anatomical connectivity of the *in vivo* human hippocampus, and track-weighted dynamic functional connectivity (TW-dFC) to investigate how direct anatomical connections between the hippocampus and the rest of the brain relate to time-varying functional interactions. In Study 1, TW-dFC maps were computed for 10 participants from the Human Connectome Project and subjected to ICA and k-means clustering to derive a data-driven parcellation of the hippocampus based on its structure-function relationships. This revealed circumscribed clusters distributed along anterior-posterior and medial-lateral axes, which broadly aligned with hippocampal subfields. In Study 2, we examined the resting-state functional connectivity profiles of each TW-dFC derived cluster in an independent sample of 100 participants. Each hippocampal cluster displayed distinct patterns of functional connectivity with specific substructures within medial temporal, parietal, frontal and occipital cortices as well as subcortical and cerebellar regions. Our findings demonstrate that TW-dFC provides a powerful framework for anatomically informed functional parcellation of the hippocampus and offers new insights into the structural-functional organisation underlying hippocampal-(sub)cortical interactions. Our approach opens new avenues for probing memory systems in health and their disruption in aging and disease.

## 1. Introduction

There is long-standing agreement that the hippocampus plays a critical role in episodic memory and supports cognitive functions such as spatial navigation (Okeefe & Nadel, 1979; Scoville & Milner, 1957). Early studies using animal models provided foundational evidence that lesions to different areas along the long (anterior-posterior) axis of the hippocampus result in deficits in specific cognitive functions (Kimura, 1958; Nadel, 1968). Building on this foundational work, recent research in humans has sought to characterise and develop a more detailed understanding of functional differences along the anterior-posterior axis of the hippocampus using *in vivo* magnetic resonance imaging (MRI) (Brunec et al., 2018; Dalton, McCormick, & Maguire, 2019; Plachti et al., 2019; Poppenk, Evensmoen, Moscovitch, & Nadel, 2013; Strange, Witter, Lein, & Moser, 2014). In parallel, increasing attention has been directed toward the putative neuroanatomical architecture that may underpin these functional differences (Dalton, D’Souza, Lv, & Calamante, 2022; Insausti & Munoz, 2001).

The hippocampus comprises cytoarchitechtonically distinct subfields (Dalton, Zeidman, Barry, Williams, & Maguire, 2017) each exhibiting unique patterns of gene expression (Strange et al., 2014), extrinsic anatomical connectivity with other parts of the brain (Insausti & Munoz, 2001), functional specialisation (Berron et al., 2016; Dalton, Zeidman, McCormick, & Maguire, 2018; Guzman, Schlogl, Frotscher, & Jonas, 2016) and functional connectivity (Dalton et al., 2019). Critically, these properties vary along their anterior-posterior axis. Functional gradients have been observed along the hippocampal long axis (Brunec et al., 2018), and, in parallel, circumscribed regions along this axis preferentially engage depending on cognitive demands (Dalton et al., 2018). Despite these increasingly detailed functional insights, a clear understanding of how functional heterogeneity along the hippocampal long axis relates to direct anatomical connections between the hippocampus and the rest of the brain remains elusive.

Structural and functional connectivity (SC and FC respectively) of the human hippocampus has been studied extensively. Notable among these studies, a method called connectivity based parcellation (CBP) has been applied to investigate hippocampal organisation as it relates to its SC (Adnan et al., 2016; Plachti et al., 2019) and FC (Chase et al., 2015; Plachti et al., 2019; Robinson et al., 2015; Robinson, Salibi, & Deshpande, 2016) with the rest of the brain. This data-driven approach has been used to parcellate the hippocampus, generating ‘maps’ of its structural and functional organisation based on connectivity profiles with the rest of the brain. These CBP derived parcellations have revealed complex organisations within the hippocampus (Adnan et al., 2016; Chase et al., 2015; Plachti et al., 2019; Robinson et al., 2015; Robinson et al., 2016). For example, Plachti and colleagues (2019) conducted CBP analysis of structural covariance (using structural covariance - CBP based on T1-weighted data) and FC (using rsFC - CBP) data to generate and compare hippocampal parcellations derived from each modality separately. Their results revealed a series of increasingly complex parcellations along the anterior-posterior and medial-lateral axes of the hippocampus for both structural covariance and FC data, ranging from simple bipartite (anterior-posterior) subdivisions to more complex organisations, depending on the clustering threshold applied, in alignment with earlier work (Adnan et al., 2016; Chase et al., 2015; Robinson et al., 2015; Robinson et al., 2016). Collectively, these studies provide important insights into SC and FC derived parcellation of the human hippocampus but, in all of them, SC and FC data have been analysed independently. Importantly, Plachti and colleages (2019) revealed a poor concordance between structural covariance and FC derived parcellation of the hippocampus, with dissimilarity being more pronounced with increased granularity (i.e., increased clustering threshold). To our knowledge, no study has integrated SC and FC data to jointly and directly investigate the relationship between SC and FC of the human hippocampus. As a result, we know surprisingly little about how differences in SC along the anterior-posterior axis of the hippocampus directly relate to its FC with other parts of the brain.

Emerging approaches to analyse multimodal MRI data now allow us to directly address this question. Advances in diffusion weighted imaging (DWI) analysis methods, provide new opportunities to map anatomical connectivity of the human hippocampus *in vivo* in unprecedented detail. For example, Dalton and colleagues (2022) recently developed and applied a novel DWI pipeline to isolate direct connections between the hippocampus and cortical mantle and map how different cortical regions anatomically connect along the anterior-posterior axis of the hippocampus using super-resolution track-density imaging (TDI: (Calamante, Tournier, Jackson, & Connelly, 2010)) and ‘endpoint density mapping’ (Dalton et al., 2022). Although the long-range cortico-hippocampal anatomical connections identified by Dalton and colleagues likely facilitate direct functional communication between the hippocampus and specific areas of the cortical mantle, no study has extended this work to investigate the relationship between these anatomical connections and their FC. One way to directly assess this is to combine this detailed DWI approach with track-weighted dynamic functional connectivity (TW-dFC (Calamante, Smith, Liang, Zalesky, & Connelly, 2017)). This approach provides a novel means to assess relationships between SC and FC by fusing SC and dynamic FC data into a quantitative 4D image (i.e., with spatial + temporal information). In simple terms, this method projects the grey matter dynamic FC information to the intersecting white matter pathways. Each track is assigned the dynamic (sliding window) Pearson’s Correlation Coefficient between the BOLD signals at its endpoints. The resulting TW-dFC maps can then be used to characterise structure-function relationships between the hippocampus and the rest of the brain. Importantly, TW-dFC differs from typical rsFC analysis. The statistical dependencies of rsFC are not limited to the underlying anatomy (Honey et al., 2009; Honey, Thivierge, & Sporns, 2010) meaning this approach is generally considered a ‘proxy’ measure of putative anatomical connectivity as it does not directly reflect anatomical connections. In contrast, TW-dFC allows us to more specifically investigate functional relationships between the hippocampus and brain areas it displays direct anatomical connections with, as characterised using our novel ‘endpoint density mapping’ approach (Dalton et al., 2022).

In this study, we aimed to systematically examine the relationship between SC and FC of the human hippocampus. We combined high-quality data from the Human Connectome Project (HCP), cutting-edge quantitative fibre-tracking methods (Calamante, 2019), a processing pipeline specifically tailored to study hippocampal SC (Dalton et al., 2022), and the TW-dFC framework (Calamante et al., 2017). To our knowledge this is the first study to merge and jointly analyse SC and FC information to investigate (sub)cortico-hippocampal interactions. We first used this data-driven approach to parcellate the human hippocampus based on its structure-function relationships with the rest of the brain (Study 1). We then used the resulting hippocampal clusters from that parcellation to analyse functional interactions between each cluster and the rest of the brain during a ‘resting state’ (Study 2). Our results represent a major advance in (i) our ability to assess relationships between SC and FC of the human hippocampus *in vivo* and (ii) our understanding of the structural-functional architecture that underpins hippocampal-dependent memory systems in the human brain.

## 2. Methods

### Study 1

In Study 1, we combined a recently developed tractography pipeline that was optimised for detailed tractography of the hippocampus (Dalton et al., 2022) with TW-dFC (Calamante et al., 2017). Independent Components Analysis (ICA) was then used to decode the TW-dFC maps and identify functional parcellations within the human hippocampus based on its direct anatomical connections and (dynamic) FC patterns with the rest of the brain.

### Participant details

Ten subjects (7 female) were selected from the minimally processed Human Connectome Project (HCP) 100 unrelated subject database (<35yo). These subjects were selected based on scan quality and visibility of the outer boundaries of the hippocampus on each participant’s T1-weighted structural MRI scan. This was done in order to increase the anatomical accuracy of our hippocampal segmentations (described below). These participants are the same participants used in our recently reported work describing the detailed anatomical connectivity of the human hippocampus (Dalton et al., 2022), and therefore the precisely delineated quantitative SC information from that study can be directly related to the findings from the current study.

### Image acquisition

The HCP diffusion protocol consisted of three diffusion-weighted shells (b-values: 1000, 2000 and 3000 s/mm^2^, with 90 diffusion-weighting directions in each shell) plus 18 reference volumes (b = 0 s/mm^2^). Each diffusion-weighted image was acquired twice, with opposite phase-encoded direction to correct for image distortion (Andersson, Skare, & Ashburner, 2003). The diffusion image matrix was 145 × 145 with 174 slices and an isotropic voxel size of 1.25 mm. The TR and TE were 5520 and 89.5 ms, respectively. Each subject also included a high resolution T1-weighted structural scan (voxel size = 0.7 mm isotropic, TR/TE = 2400/2.14 ms, flip angle = 8°) and a high resolution T2-weighted structural scan (voxel size = 0.7 mm isotropic, TR/TE= 3200/565 ms and variable flip angle).

The resting-state functional magnetic resonance imaging (rsfMRI) data were collected using Gradient-echo EPI, with TR=0.72s, TE=33.1ms, flip angle of 52 degrees, FOV= 208×208 mm^2^, 2 mm isotropic resolution. A multiband factor of 8 was used to accelerate the scan. During the 14.5 minute scan, 1200 volumes were collected (S. M. Smith et al., 2013). The rsfMRI is minimally processed and projected to the new data representation of cortical surface+ subcortical volume, i.e., in the CIFTI format (M. F. Glasser et al., 2013). These data were used for both Study 1 and Study 2 (described below).

### Manual segmentation of the hippocampus

The whole hippocampus was manually segmented for each participant on coronal slices of the T1-weighted image using ITK-SNAP (Yushkevich et al., 2006). Considering automated methods of hippocampal segmentation are sometimes not sufficiently accurate, particularly in the anterior and posterior most extents of the hippocampus (see Dalton et al., 2022 for representative examples), manual segmentation was essential for this detailed work. We manually segmented the whole hippocampus of each participant following procedures described previously (Dalton et al., 2022).

### Image pre-processing

All DWI analyses described here were conducted as part of our previously reported work (Dalton et al., 2022) where the interested reader can find a detailed description. In brief, the HCP minimally processed DWI data was further pre-processed using the MRtrix software package (http://www.mrtrix.org) (J.-D. Tournier et al., 2019; J. D. Tournier, Calamante, & Connelly, 2012). Additional processing steps were implemented in accordance with previous work (Civier, Smith, Yeh, Connelly, & Calamante, 2019) and included bias-field correction (Tustison et al., 2010) as well as multi-shell multi-tissue constrained spherical deconvolution to generate a fibre orientation distribution (FOD) image (Jeurissen, Tournier, Dhollander, Connelly, & Sijbers, 2014; J.-D. Tournier, Calamante, Gadian, & Connelly, 2004; J. Donald Tournier, Fernando Calamante, & Alan Connelly, 2007). The T1-weighted image was used to generate a ‘five-tissue-type’ (5TT) image (R. E. Smith, Tournier, Calamante, & Connelly, 2012) using FSL (Patenaude, Smith, Kennedy, & Jenkinson, 2011; S. M. Smith, 2002; S. M. Smith et al., 2004; Zhang, Brady, & Smith, 2001): tissue 1 = cortical grey matter, tissue 2 = sub-cortical grey matter, tissue 3 = white matter, tissue 4 = CSF, and tissue 5 = pathological tissue (For healthy subjects, tissue 5 is typically an empty image).

Besides the steps carried out by the HCP team as part of the minimally processed fMRI datasets, we performed additional pre-processing steps to enhance signal quality and reduce noise in both spatial and temporal dimensions. Spatial smoothing was performed using a Gaussian filter with a full-width at half-maximum (FWHM) of 6mm. Following this, a bandpass filter was applied to maintain frequencies within the range of 0.01 to 0.1Hz, effectively isolating the desired signal bandwidth.

### Whole-brain tractogram

As described in (Dalton et al., 2022), the FOD data (Jeurissen et al., 2014; J. D. Tournier, F. Calamante, & A. Connelly, 2007) and the 5TT image were used to generate 70 million anatomically constrained tracks (R. E. Smith et al., 2012) using dynamic seeding (R. E. Smith, Tournier, Calamante, & Connelly, 2015b) and the 2nd-order Integration over Fibre Orientation Distributions (iFOD2 (J Donald Tournier, Calamante, & Connelly, 2010)) probabilistic fibre-tracking algorithm (see (Dalton et al., 2022) for tracking algorithm parameters).

### Hippocampus tractogram

We recently developed a tailored pipeline that enables researchers to track streamlines into the hippocampus, identify the location of their termination points with high spatial precision and isolate only streamlines with an endpoint in the hippocampus. The interested reader can find a detailed description of this novel approach in (Dalton et al., 2022). Using this approach, we generated an additional 10 million anatomically constrained tracks (R. E. Smith et al., 2012) seeded from the manually segmented hippocampus. The 70 million whole-brain tracks and the 10 million hippocampus tracks were combined, and Spherical-deconvolution Informed Filtering of Tractograms 2 (SIFT2 (R. E. Smith et al., 2015b)) was used on the combined 80 million track file, thereby assigning a weight contribution to each track for improved biological credence to the connectivity measurements (R. E. Smith, Tournier, Calamante, & Connelly, 2015a). Tracks (and SIFT2 weights) which had an endpoint in the hippocampus were extracted (referred to here as the ‘hippocampus tractogram’) and used in the subsequent steps.

The power of this approach in the context of the current study, is that it allowed us to identify and isolate only streamlines with a well-defined endpoint in the hippocampus, thereby restricting our subsequent TW-dFC analyses to focus specifically on direct connections between the hippocampus and the rest of the brain.

### Image registration into the template space

All subsequent analyses were conducted in a common template space. Given that both accurate registration of white matter (for fibre-tracking) and grey matter (for rsfMRI) is critical, we used our recently developed multi-modal tissue-unbiased template (Lv, Zeng, Ho, D’Souza, & Calamante, 2023) to increase the reliability of our results. The template in (Lv et al., 2023) was constructed by jointly considering T1w, T2w, as well as fractional anisotropy and mean diffusivity maps computed from a tensor modelling of DWI data (Basser & Pierpaoli, 2011) - see (Lv et al., 2023) for further details. The multimodal MRI data of each individual is registered to the template space and the derived transformation was applied to the fibre-tracks (hippocampus tractogram) and rsfMRI data of each individual.

### TW-dFC Analysis

We calculated the TW-dFC map for the hippocampus tractogram by assigning each streamline a ‘dynamic functional weighting’ given by the (sliding window) functional correlation between rsfMRI data at its endpoints (Calamante, 2017; Calamante et al., 2017). Each streamline of the hippocampus tractogram has one endpoint in the hippocampus and another in either the cortical mantle or a subcortical structure, and each streamline is assigned the (dynamic) FC between its endpoint in the hippocampus and its associated endpoint in the (sub)cortex (see Figure 1 for a schematic representation). The (dynamic) FC value is further mapped to the voxels that each streamline traversed. When multiple streamlines traverse the same voxel, the (dynamic) FC is averaged across them (Figure 1). In essence, this method projects grey matter (dynamic) FC information to the intersecting white matter pathways. Based on previous work (Calamante et al., 2017), TW-dFC maps were computed in the multimodal template space with the Hamming window size of 85 time points (corresponding to a sliding window of 61.2 sec) at 2mm isotropic resolution, and the resulting 4D TW-dFC data were subsequently analysed using ICA (described below).

**Figure 1.**
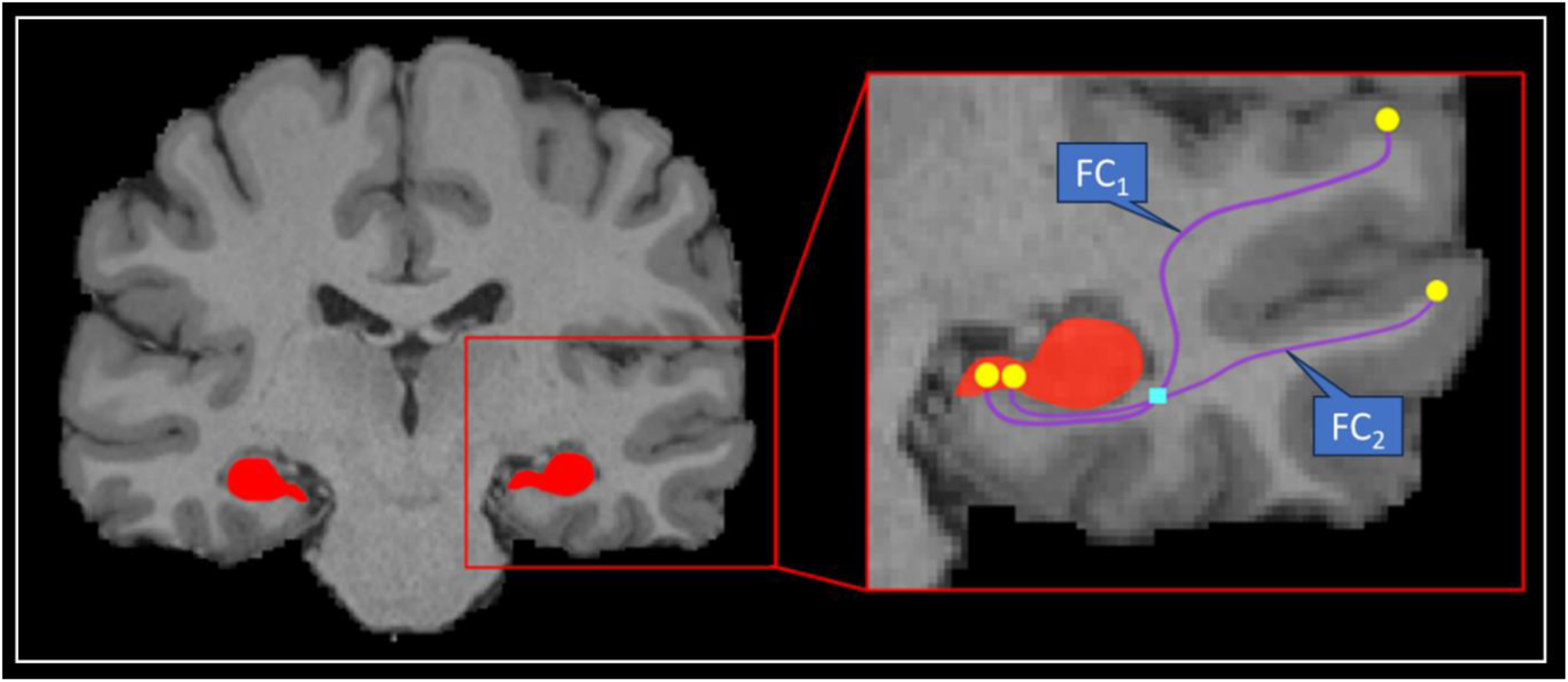
Schematic Representation of TW-dFC. Schematic example of assignment of (dynamic) functional connectivity (FC) weighting to each track (indicated by the blue FC_i_ label associated with each track). In this example, two streamlines are represented (purple lines), each with an endpoint in the hippocampus and cortical ribbon (yellow circles). Each track is assigned the temporal Pearson correlation between the BOLD signals at its endpoints (yellow circles). For any given voxel (e.g., cyan square), the TW-dFC intensity in that voxel corresponds to the average of the weightings of all tracks traversing the voxel: (FC_1_ + FC_2_ /2).

##Supplementary Video 1 presents a representative TW-dFC map from a single participant and offers a striking visualisation of time-varying hippocampal connectivity. This dynamic mapping highlights how the hippocampus functionally interacts with distinct (sub)cortical regions over time, constrained by its anatomical connections. Functional coupling with the rest of the brain is projected onto the white matter pathways that mediate these interactions, revealing how specific tracts support communication between the hippocampus and distributed brain networks throughout the resting-state scan. We discuss this further in the Discussion section.

### Independent Components Analysis (ICA)

The TW-dFC data of 10 subjects was temporally concatenated and decomposed into 180 independent components with the Melodic-ICA toolbox from FSL (S. M. Smith et al., 2004). The number of components was chosen to make sure the ICA components explain >98% of the variance of the data while avoiding redundancy. Each ICA component, is represented by a spatial z-score map indicating the component’s weight on each voxel.

### Clustering

An additional whole hippocampus *group* mask was manually delineated on the template level T1-weighted image in accordance with previous work (Dalton et al., 2022). The ICA results (i.e. the set of spatial maps) were used to identify clusters within the hippocampus by thresholding the z-score maps at p>0.5. In essence, this allowed us to characterise, in a data-driven manner, spatially distinct (SC-driven) functional clusters within the hippocampus. We used the group level hippocampus mask to extract the z-scores of 180 ICA components at each voxel within the hippocampus. Therefore, we formed a 180-dimension feature vector for each hippocampal voxel, with which we clustered the whole hippocampus into functional clusters using the k-mean clustering method. We generated results for k = 4-, 6- and 8-cluster solutions. For simplicity, we present detailed results for the 6-cluster solution in the main text, as this provides a reasonable compromise between level of clustering subdivision and ease of interpretation of the resulting networks. We provide a visual representation of the 4-, 6- and 8-cluster solutions in Supplementary Figure 1. A more detailed representation of the 6-cluser solution is provided in Figure 2 and described in the Results section.

**Figure 2.**
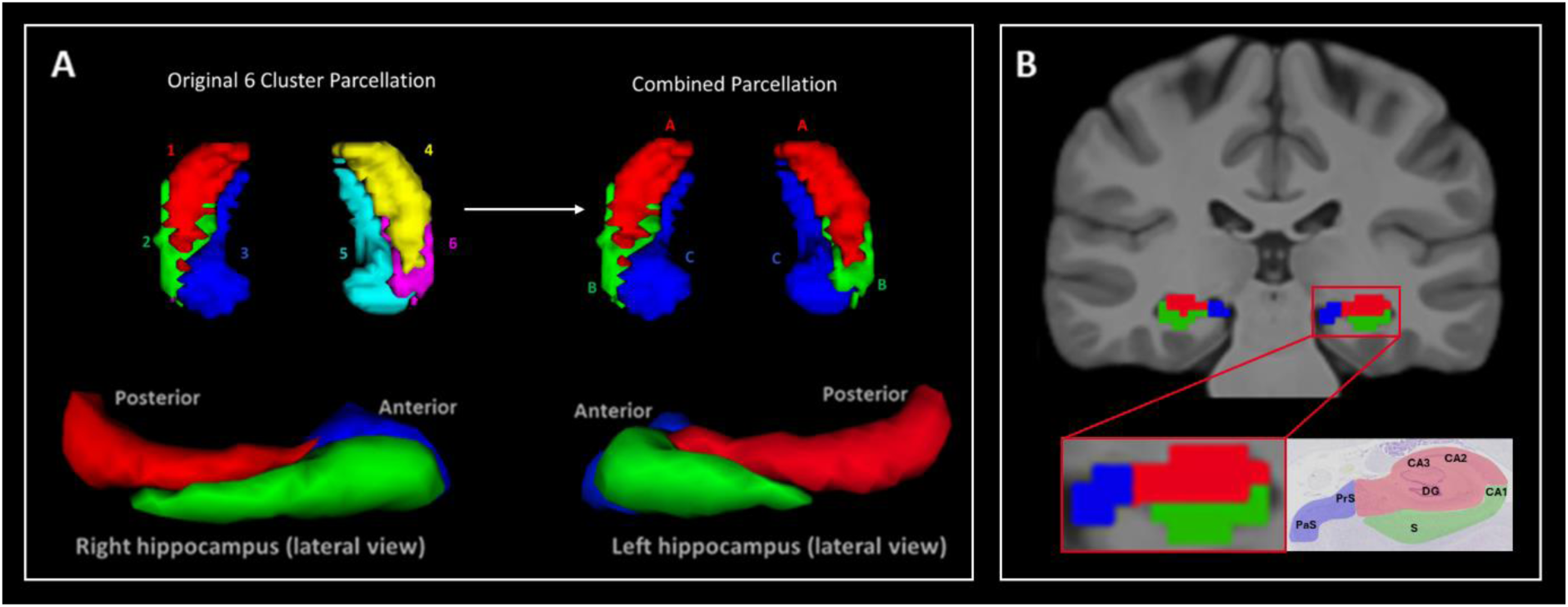
Results of the 6 cluster ICA analysis. (A) 3D rendering of clusters revealed by k-means clustering of ICA of TW-dFC data distributed along the anterior-posterior and medial-lateral axes of the hippocampus. The original TW-dFC derived 6-cluster parcellation from Study 1 is presented (top left) along with the combined parcellation (top right) which combined equivalent clusters in the left and right hippocampus into single bilateral clusters (Clusters A, B and C), used for rsfMRI analysis in Study 2. The amended parcellation is displayed viewed from a superior view of the bilateral hippocampus (top right) and lateral views of the right (bottom left) and left (bottom right) hippocampus. Note the clear overlapping nature of clusters A and B when viewed from the lateral perspective. (B) Representative example of TW-dFC derived clusters overlaid on a T1-weighted image in the coronal plane (top). Note each cluster roughly aligns with the location of cytoarchitecturally defined hippocampal subfields. This is clear when comparing the location of each cluster with anatomically defined subfields on histology (bottom images). For ease of visualisation, we have overlaid a rough estimate schematic representation of the TW-dFC functional clusters over a histological image (bottom right). Note, Cluster 1 (red) roughly encompasses the location of DG/CA3/CA2. Cluster 2 (green) roughly encompasses the location of CA1 and subiculum (S). Cluster 3 (blue) roughly encompasses the location of the presubiculum (PrS) and parasubiculum (PaS).

### Study 2

In the complementary Study 2, based on the hippocampus clustering results generated using the joint structural and (dynamic) functional connectivity analysis from Study 1, we conducted a subsequent series of seed-based FC analyses to identify the distinct functional networks associated with each hippocampal cluster.

### Participant details

As Study 2 does not require the definition of subject-specific hippocampal masks (which are time consuming due to the nature of manual delineation), a much larger subject group was included to increase the reliability of the identified functional networks associated with each hippocampal cluster. Thus, Study 2 involved data from one hundred subjects (54 female; <35yo, mean age = 29.1 ± 3.7) available in the minimally processed Human Connectome Project (HCP) 100 unrelated subject database (Van Essen et al., 2013).

### Image acquisition and pre-processing

Image acquisition and preprocessing steps are described in the ‘Image Acquisition’ and ‘Image preprocessing’ sections in Study 1.

### Seed-based Functional Connectivity Calculation

We used the pre-processed CIFTI rsfMRI data and the hippocampal TW-dFC cluster atlas derived from Study 1 to calculate the seed-based FC (Fox et al., 2005) between each hippocampal cluster and the rest of the brain. The results of Study 1 revealed clusters in the left and right hippocampus that were strikingly equivalent in their symmetry (described in the Results section). We therefore combined equivalent clusters from the left and right hippocampus into a single mask, reducing the initial six clusters to three for the computation of FC networks in Study 2. Specifically, clusters 1 and 4 were merged to make bilateral cluster A, clusters 2 and 6 were combined to make bilateral cluster B and clusters 3 and 5 were merged to make bilateral cluster C (see Figure 2A for visualisation). Each of the resulting bilateral clusters served as seeds to calculate their associated FC map.

For each cluster, we first extracted the time series data of all voxels within the cluster mask. The time series associated with the cluster were then averaged to create the representative time series for each cluster. This averaged time series was subsequently used to calculate the seed-based FC for that cluster. We employed Pearson’s correlation to measure the FC between the averaged time series of each cluster and the time series of the whole brain.

### T-Tests

Following the calculation of seed-based FC of each hippocampal cluster, we conducted comparative analyses of their FC. To achieve this, we employed one-way t-tests to evaluate differences between seed FC maps. This analysis comprised six distinct t-tests which aimed to assess how patterns of preferential FC between each cluster and the rest of the brain differs compared to the other clusters. Specifically, using this approach, we identified (sub)cortical regions that; cluster A has greater FC with relative to cluster B (A > B) and vice versa (B > A), that cluster A exhibits greater FC with relative to cluster C (A > C) and vice versa (C > A), and that cluster B has greater FC with relative to cluster C (B > C) and vice versa (C > B). This comprehensive set of tests was designed to thoroughly explore and elucidate the relative functional connectivity strengths among the clusters. P-values obtained from these tests were corrected through false discovery rate (FDR) correction, after which, they were converted to z-scores. We mapped each z-score to the brain surface using a threshold of 1.645 (equivalent to p=0.05).

### Cortical parcellation

FreeSurfer (Fischl, 2012) was used to process the group average T1-weighted image. The Human Connectome Project Multi-Modal Parcellation 1.0 (HCPMMP) (Matthew F. Glasser et al., 2016) was mapped to the group averaged T1w image in accordance with previous work (Tahedl, 2020). The parcellation divided the cerebral cortex into 360 parcels (180 per hemisphere). Results relating to FC between the hippocampus and cortical mantle are reported and interpreted according to HCPMMP delineations.

## 3. Results

### Study 1

Our method was effective in identifying differentiable functional clusters within the human hippocampus based on combined structural and (dynamic) function connectivity. The results of k-means clustering of ICA of TW-dFC maps revealed 6 functional clusters distributed along the anterior-posterior and medial-lateral axes of the hippocampus (see Figure 2A). The left and right hippocampus each contained 3 clusters that were strikingly symmetrical across the left and right hippocampus, despite no such constraint having been imposed. These consisted of the following; a cluster that encompassed the entire hippocampal tail but extended anteriorly, specifically encompassing superior portions of the hippocampal body, roughly aligning with the DG/CA3/CA2 subfields (See Figure 2A and B; Cluster A - red); A lateral cluster, encompassing inferolateral portions of the head and body of the hippocampus, roughly aligning with CA1 and subiculum subfields (See Figure 2A and B; Cluster B - green); A medial cluster encompassing hippocampal subfields within the uncus and extending posteriorly along the medial edge of the hippocampus, roughly aligning with the location of the distal subiculum, presubiculum and parasubiculum (See Figure 2A and B; Cluster C - blue). The alignment of functional clusters with these subfields was primarily observed in the body of the hippocampus. Functional clusters in the hippocampal head and tail more broadly encompassed multiple subfields suggesting coarser functional organisation in these regions (see Supplementary Figure 2 for representative examples of the entire anterior-posterior extent of each TW-dFC derived cluster).

It should be noted that although these clusters were generated using an entirely data-driven approach, the k-means clustering method requires to specify the total number of clusters to be generated. As described in the Methods section, we focus on the 6-cluster solution here but provide a visual representation of the 4- and 8-cluster solutions in Supplementary Figure 1. The TW-dFC derived functional parcellations generated in this study have been made publicly available to the research community through OSF (To be uploaded on acceptance).

### Study 2

Group level seed-based rsfMRI analysis revealed that each of the TW-dFC-guided hippocampal clusters were functionally associated with distinct cortical and subcortical regions. For brevity, we illustrate these differences by focussing on results from 3 t-test comparisons: cluster A > cluster C; cluster C > cluster A and; cluster B > cluster C (see Figure 3 for representative examples). The interested reader can find results relating to all contrasts in Supplementary Table 1 and a more detailed, labelled visualisation of results for all contrasts are provided in Supplementary Figures 3-8.

**Figure 3.**
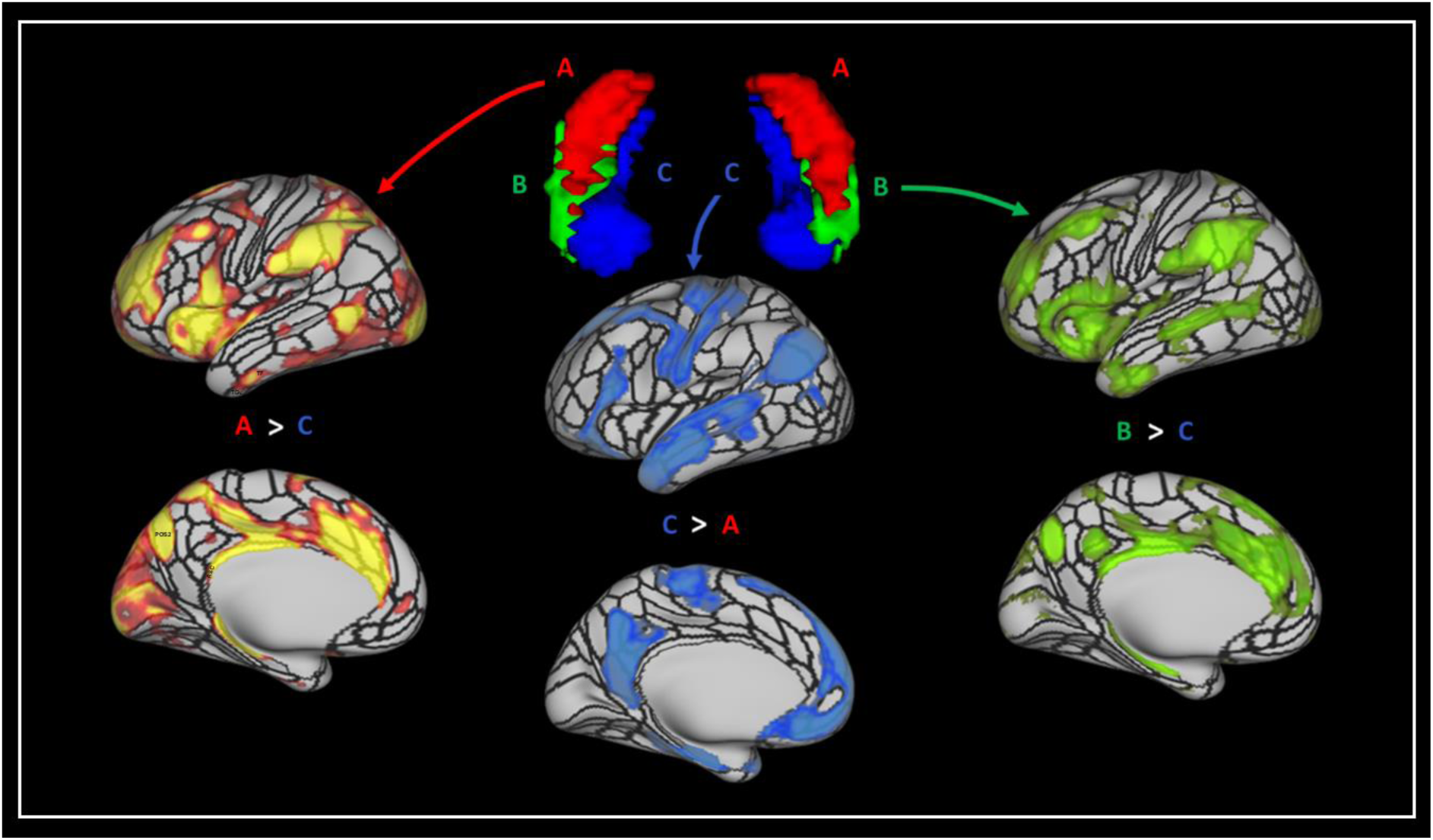
Results of the seed-based rsfMRI analysis based on the TW-dFC hippocampal clusters. Left - Results for the contrast of cluster A (hippocampal tail cluster) > cluster C (medial cluster); Middle - cluster C (medial cluster) > cluster A (hippocampal tail cluster) and; Right - cluster B (anterolateral cluster) > cluster C (medial cluster). T-test results are thresholded at p<0.05 FDR corrected.

The contrast of cluster A (hippocampal tail cluster) > cluster C (medial cluster) (See Figure 3 and Supplementary Figure 4 for a more detailed and labelled visualisation) revealed that compared to the medial cluster, the hippocampal tail cluster showed preferential functional connectivity with primary and early visual (V1, V2, V3, V4); dorsal stream visual (V6, V6A, V3B); ventral stream visual (VMV3, PIT); visual areas neighbouring the MT+ complex (PH, LO1, LO2, LO3, V3CD); paracentral lobular and mid cingulate cortex (6ma, SCEF, 24dv, 5mv); premotor cortex (FEF, PEF, 6r), posterior opercular cortex (43, PFcm); auditory association cortex (STSdp, STSvp); insular and frontal opercular cortex (PI, PoI1, PoI2, FOP3, FOP4, FOP5, MI, AVI, AAIC, Pir); medial temporal cortex (peEC); lateral temporal cortex (PHT, TE2p, TE2a, TE1m); temporo-parieto-occipital junction (TPOJ2, PSL); superior parietal cortex (7AM, 7PL, 7AL, AIP, LIPd, LIPv, MIP); inferior parietal cortex (PFOP, PF, PFm, IP1, IP2, pGp); posterior cingulate cortex (RSC, 23c, 23d, 31pv, 31pd, POS2, DVT); anterior cingulate and medial prefrontal cortex (p24pr, a24pr, p32pr, a32pr, 8BM, p24, p32, d32); orbital and polar frontal cortex (pOFC, 13l, 11l, a10p, p10p); inferior frontal cortex (p47r, IFSa, 44) and dorsolateral prefrontal cortex (a9- 46v, 9-46d, 46, P9-46v, 8C). Compared to cluster C, cluster A also displayed preferential connectivity with subcortical structures including thalamus, putamen, pallidum, caudate nucleus, and with specific cerebellar lobules.

The contrast of cluster C (medial cluster) > cluster A (hippocampal tail cluster) (See Figure 3 and Supplementary Figure 7 for a more detailed and labelled visualisation) revealed that compared to the hippocampal tail cluster, the medial cluster showed preferential functional connectivity with ventral stream visual (FFC); MT+ complex (MT); somatosensory and motor cortex (1, 2, 3a, 3b, 4); paracentral lobular and mid cingulate cortex (24dd, 5m, 5L); premotor cortex (6v, 6d, 55b), auditory association cortex (A4, A5, STSdp, STSda, STSva, STGa); medial temporal cortex (EC, PHA1, PHA2, PHA3); lateral temporal cortex (TE1p, TE1a, TE2a, TGd); temporo-parieto-occipital junction (TPOJ1, TPOJ3, STV); superior parietal cortex (7Pm); inferior parietal cortex (PGi, PGs); posterior cingulate cortex (d23ab, v23ab, 7m, POS1, DVT, ProS); anterior cingulate and medial prefrontal cortex (25, s32, 10r, 10v, 9m, a24); orbital and polar frontal cortex (10d, 9a, 47m); inferior frontal cortex (47l, 45, IFSp, IFJp) and dorsolateral prefrontal cortex (8C, 8AV, 8AD). Compared to cluster A, cluster B also displayed preferential connectivity with subcortical structures, specifically with the amygdala and with specific cerebellar lobules.

The contrast of cluster B (lateral cluster) > cluster C (medial cluster) (See Figure 3 and Supplementary Figure 6 for a more detailed and labelled visualisation) revealed that compared to the medial cluster, the lateral cluster showed preferential functional connectivity with primary and early visual (V1, V3, V4); dorsal stream visual (V6); paracentral lobular and mid cingulate cortex (6ma, SCEF, 24dv, 5mv); premotor cortex (PEF, 6r, 55b), posterior opercular cortex (43, PFcm); auditory association cortex (STSdp, STSvp); insular and frontal opercular cortex (PI, PoI1, PoI2, FOP3, FOP4, FOP5, MI, AVI, AAIC); medial temporal cortex (peEC); lateral temporal cortex (PHT, TE1m, TE2p, TE2a, TGv, TGd); temporo-parieto-occipital junction (TPOJ1, TPOJ2, PSL); superior parietal cortex (AIP, 7Am); inferior parietal cortex (PFOP, PF, PFm, IP2, PGi, PGs); posterior cingulate cortex (RSC, d23ab, 31pv, 31pd, 23c, 23d, POS2); anterior cingulate and medial prefrontal cortex (p24pr, a24pr, p32pr, a32pr, 8BM, p24, d32, p32, 9m); orbital and polar frontal cortex (47s, 11l, a10p); inferior frontal cortex (47l, 45, 44, p47r, IFSa) and dorsolateral prefrontal cortex (a9-46v, 9-46d, 9a, 9p, 46, 8C, 8AV, P9-46v). Compared to cluster C, cluster B also displayed preferential connectivity with subcortical structures including caudate nucleus and, to a lesser degree, with the putamen, pallidum and thalamus, and with specific cerebellar lobules.

## 4. Discussion

Advancing our knowledge of how the hippocampus is structurally and functionally connected to the broader brain may be key to deciphering the principles that govern functional differentiation along the hippocampal long axis. Having previously demonstrated our ability to create detailed maps of SC between the hippocampus and cortical mantle in the *in vivo* human brain (Dalton et al., 2022), here we extended this work by examining the relationship between SC and rsFC in young healthy adults. We leveraged TW-dFC to integrate structural and functional data and probe structure-function associations between the hippocampus and brain areas with which it displays direct anatomical connections. Through this innovative approach, we identified distinct functional clusters along the anterior-posterior and medial-lateral axes of the hippocampus that each exhibit unique patterns of structural-guided (dynamic) functional connectivity with specific cortical and subcortical areas. Our results represent a significant advance in understanding how anatomical connectivity between the human hippocampus and the broader brain supports its internal functional differentiation and shed new light on the intrinsic organisational principles of the hippocampus itself. Our results suggest that functional connectivity of the hippocampus may follow distinct, spatially structured patterns that reflect both its internal architecture and its anatomical connectivity with the rest of the brain. Our method provides a powerful new framework to interrogate structure-function relationships between the hippocampus and its distributed networks in health and offers a foundation for future research into how age-related or disease-associated disruptions to these anatomical circuits may contribute to memory decline, particularly in conditions such as Alzheimer’s disease.

### 4.1 Anatomically constrained functional parcellation of the human hippocampus

This study introduces a novel application of TW-dFC to map time-resolved functional interactions between the hippocampus and brain regions with which it shares direct anatomical connections. Building on a DWI pipeline we previously developed to delineate hippocampal white matter pathways with high spatial precision (Dalton et al., 2022), we first identified cortical and subcortical voxels that exhibit direct SC with the hippocampus. We then extended this anatomically grounded framework by projecting dynamic functional connectivity estimates from grey matter onto these structurally defined pathways, thereby isolating time-varying fluctuations in hippocampal functional coupling constrained by direct anatomical connections (Figure 1). This approach provides a key advantage over conventional methods by enabling precise targeted analysis of dynamic connectivity between the hippocampus and brain voxels with which it displays direct structural connections, eschewing indirect or network-level associations in favour of a fine-grained, anatomically constrained mapping. Supplementary Video 1 is compelling in that it visualises dynamic fluctuations in how the hippocampus functionally interacts with different brain regions over the time-course of the resting-state functional scan, projected onto the white matter pathways that mediate these interactions. This offers a vivid depiction of how hippocampal connectivity shifts across distributed brain regions over time. To our knowledge, this is the first study to directly visualise, quantify, and track dynamic functional connectivity between the hippocampus and its structurally connected targets. When combined with independent component analysis (ICA) and k-means clustering, this framework revealed intricate patterns of functional clustering organised along both the anterior-posterior and medial-lateral axes of the hippocampus. Together, these findings provide a data-driven window into the organisation of hippocampal connectivity dynamics with the rest of the brain during a ‘resting state’ and lay the foundation for future investigations into how these patterns relate to cognition and behaviour.

For simplicity, we focus on the 6-cluster solution described in the results section. This solution yielded three clusters within the hippocampus in each hemisphere, with a high degree of spatial symmetry across hemispheres. We combined the equivalent left and right hippocampal clusters into three bilateral clusters referred to as clusters A, B and C (described in detail in Results; see Figure 2). Clusters A and B broadly correspond to the hippocampal tail and anterior-lateral portions of the hippocampus, respectively. However, rather than a clear anterior-posterior division, these clusters exhibited substantial topographic overlap within the hippocampal body (Figure 2A), where each aligned preferentially with cytoarchitectonically defined subfields (Figure 2B; see Results for details). It is quite compelling that, although these clusters were derived in a fully data-driven manner based on anatomically constrained dynamic functional connectivity patterns, their topography within the hippocampal body closely mirrored known subfield architecture. In contrast, cluster C encompassed uncal portions of the hippocampal head and extended posteriorly to encompass medial portions of the hippocampus along almost its entire anterior-posterior extent (see Figure 2A and Supplementary Figure 2; see Results for details). These findings challenge the utility of simplified anterior-posterior models of hippocampal organisation, highlighting instead a more nuanced internal structure defined by overlapping spatial gradients and subfield-specific functional integration. We now discuss how our TW-dFC derived parcellations relate to the extant literature on structural and functional connectivity based hippocampal parcellation.

### 4.2 Using TW-dFC to bridge the gap between SC and FC

Several studies have applied CBP approaches to conduct hippocampal parcellation based on structural or functional connectivity, with results ranging from simple bipartite anterior-posterior subdivisions to more complex organisations (Adnan et al., 2016; Chase et al., 2015; Plachti et al., 2019; Robinson et al., 2015; Robinson et al., 2016). Consistent with these studies, our results reflected an increasing parcellation complexity along both anterior-posterior and medial-lateral axes with higher clustering thresholds (see Supplementary Figure 1). Perhaps most relevant to our study, Plachti and colleagues (2019) compared parcellations derived from structural covariance and rsFC using CBP methods. They reported poor concordance between structural covariance and rsFC parcellations, with increasing dissimilarity between modalities at higher clustering thresholds, underscoring the complex structure-function relationships that remain incompletely understood. By combining SC and FC data, our results bridge this gap by leveraging the complementary information provided by each modality and strike a balance between previously reported structural covariance and rsFC-based parcellations.

For comparison, we examine our findings alongside those of Plachti and colleagues (2019). In their 3-cluster solution (equivalent to our 6-cluster solution; i.e., 3 clusters per hippocampus), Plachti and colleagues reported similar but distinct patterns across structural covariance and rsFC derived parcellations. In brief, structural covariance parcellation revealed an anterior cluster encompassing the hippocampal head (see Figure 4A; left, green), a posterior lateral cluster encompassing the posterior lateral and tail portions of the hippocampus (Figure 4A; left, red), and a more circumscribed medial intermediate cluster (Figure 4A; left, blue). In contrast, their rsFC-derived parcellation identified an anteromedial cluster encompassing the medial part of the hippocampal head (Figure 4A; right, blue), an anterolateral cluster encompassing lateral portions of the head and intermediate body (Figure 4A; right, green), and a posterior cluster encompassing the posterior body and hippocampal tail (Figure 4A; right, red). Our TW-dFC-derived parcellation recapitulates aspects of these structural covariance and rsFC derived clustering patterns but extends this work by providing a parcellation derived from a combined structure-function analysis.

**Figure 4.**
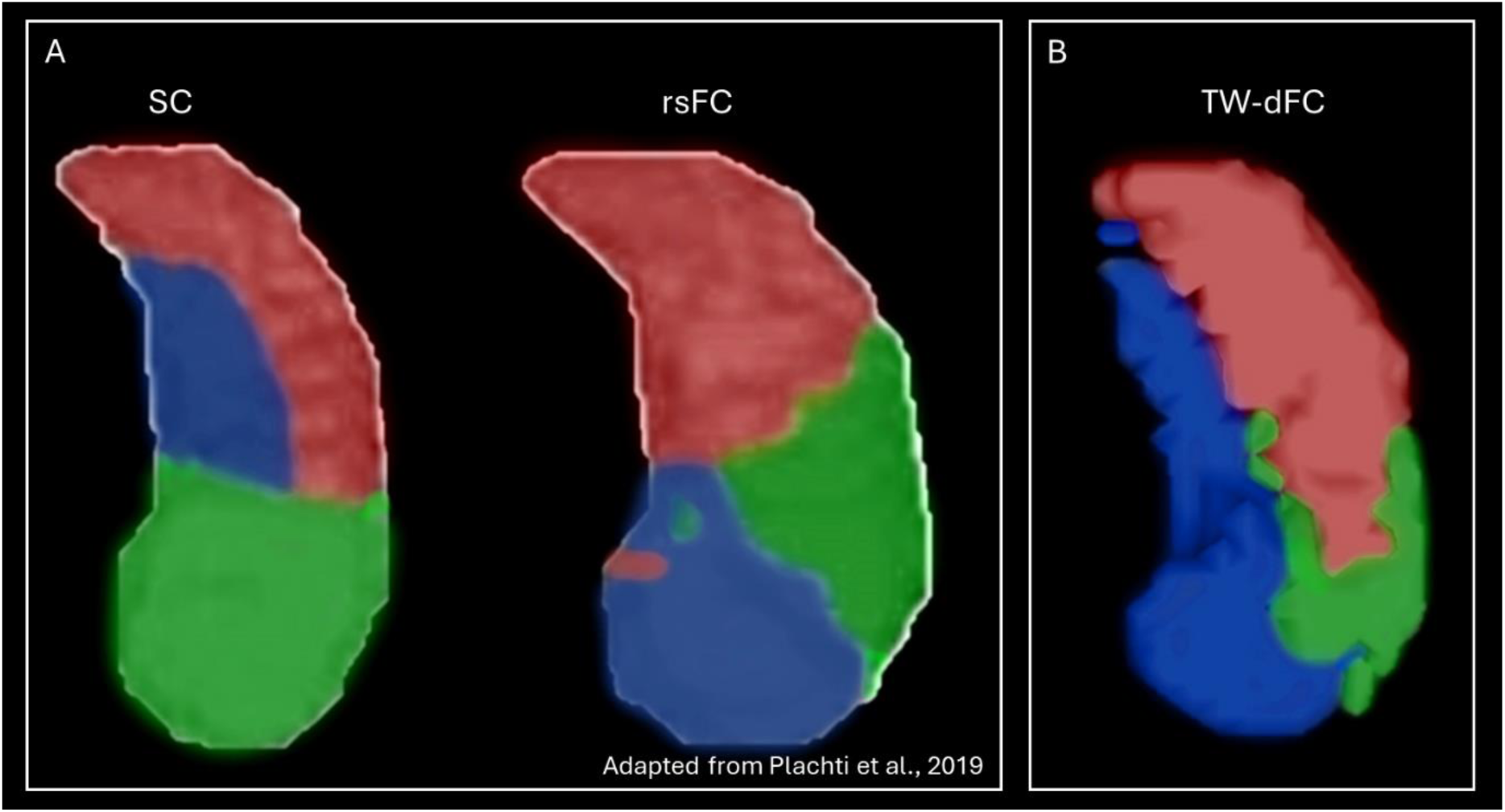
Comparison of structural covariance, rsFC and TW-dFC parcellation of the hippocampus. 3D rendering of structural covariance (A; left) and rsFC (A; right) derived clusters described by Plachti and colleagues (2019)* and the TW-dFC derived clusters reported in the current study (B). Note the TW-dFC derived clusters strike an interesting middle ground between the previously reported structural covariance and rsFC derived parcellations presented in A. *For ease of comparison, we have created recoloured representations of Plachti and colleagues (2019) findings to more closely match the colour scheme used in the current study.

Specifically, Cluster A in our study (Figure 4B; red) resembles Plachti and colleagues structural covariance-derived posterior cluster (Figure 4A; left, red) but extends further anteriorly and overlies Cluster B. Cluster B (Figure 4B; green) aligns with Plachti and colleagues rsFC-derived anterolateral cluster (Figure 4A; right, green) but extends more posteriorly and inferior to Cluster A. Intriguingly, our TW-dFC derived Cluster C (Figure 4B; blue) resembles a combination of Plachti and colleagues structural covariance derived intermediate medial cluster (Figure 4A; left, blue) and rsFC-derived anteromedial cluster (Figure 4A; right, blue) and strikes an interesting middle ground between these prior data-driven parcellations. Overall, the results of Study 1 demonstrate that TW-dFC is a viable method to integrate quantitative measures of anatomical connectivity strength with dynamic FC to bridge discrepancies previously observed between SC and FC parcellation. Indeed, it must be emphasised that, in contrast to previous approaches, our parcellation is derived from dynamic functional connectivity between only those (sub)cortical voxels that display direct anatomical connections with the hippocampus (isolated using our novel DWI pipeline (Dalton et al., 2022)) and must be interpreted with this in mind. This innovative approach provides fresh insights into the structural-functional organisation of the hippocampus as it relates to directly connected brain areas. We now briefly discuss functional associations between each of the TW-dFC derived hippocampal clusters and the rest of the brain observed in Study 2.

### 4.3 TW-dFC derived hippocampal clusters display unique patterns of functional connectivity

In Study 2, we used the hippocampal clusters derived from the TW-dFC parcellation as seed regions for rsFC analyses to map the distributed functional networks associated with each cluster. Our findings revealed intricate patterns of FC between individual hippocampal clusters and specific cortical, subcortical, and cerebellar regions. Overall, the observed connectivity profiles were broadly consistent with well-established functional associations between, for example, the anterior hippocampus and temporal/vmPFC regions and between the posterior hippocampus and parietal/occipital regions (Adnan et al., 2016; Barnett et al., 2021; Dalton et al., 2019; Knapen, 2021; Poppenk & Moscovitch, 2011; Tang et al., 2020). However, the anatomical specificity of certain associations are particularly striking and suggest that the distinct functional subunits within the hippocampus, identified by our TW-dFC parcellation, each preferentially interact with discrete subcomponents within established brain networks. We highlight some key observations below.

Different TW-dFC derived hippocampal clusters showed distinct patterns of functional connectivity with specific subregions within the PCC, revealing a fine-grained topography through which discrete hippocampal zones may selectively engage with distinct components of the PCC. For example, Cluster A showed preferential FC specifically with dorsal portions of the RSC and POS2 (see Figure 5, top left and Supplementary Figures 3 and 4), while Clusters B and C were more broadly associated with ventral RSC and areas POS1, v23ab, d23ab, 31pv, 31pd and 7m within the PCC (see Figure 5, top right and Supplementary Figures 5 and 7). While previous research has established anatomical (Dalton et al., 2022; Kravitz, Saleem, Baker, & Mishkin, 2011) and functional (Dalton et al., 2018; Leech & Smallwood, 2019) links between the hippocampus and specific PCC subregions, the close proximity of PCC subregions has traditionally hindered clear separation of their functional signals due to insufficient resolution in human neuroimaging studies. Considering different subregions within both the hippocampus and PCC are implicated in different elements of visuospatial processing and related cognition (Dalton & Maguire, 2017; Epstein, 2008; Silson, Steel, & Baker, 2016), our TW-dFC derived hippocampal parcellation may provide a powerful tool for researchers seeking to unravel fine-grained, anatomically grounded interactions between the hippocampus and distinct subregions within the PCC during visuospatial and related cognitive tasks.

**Figure 5.**
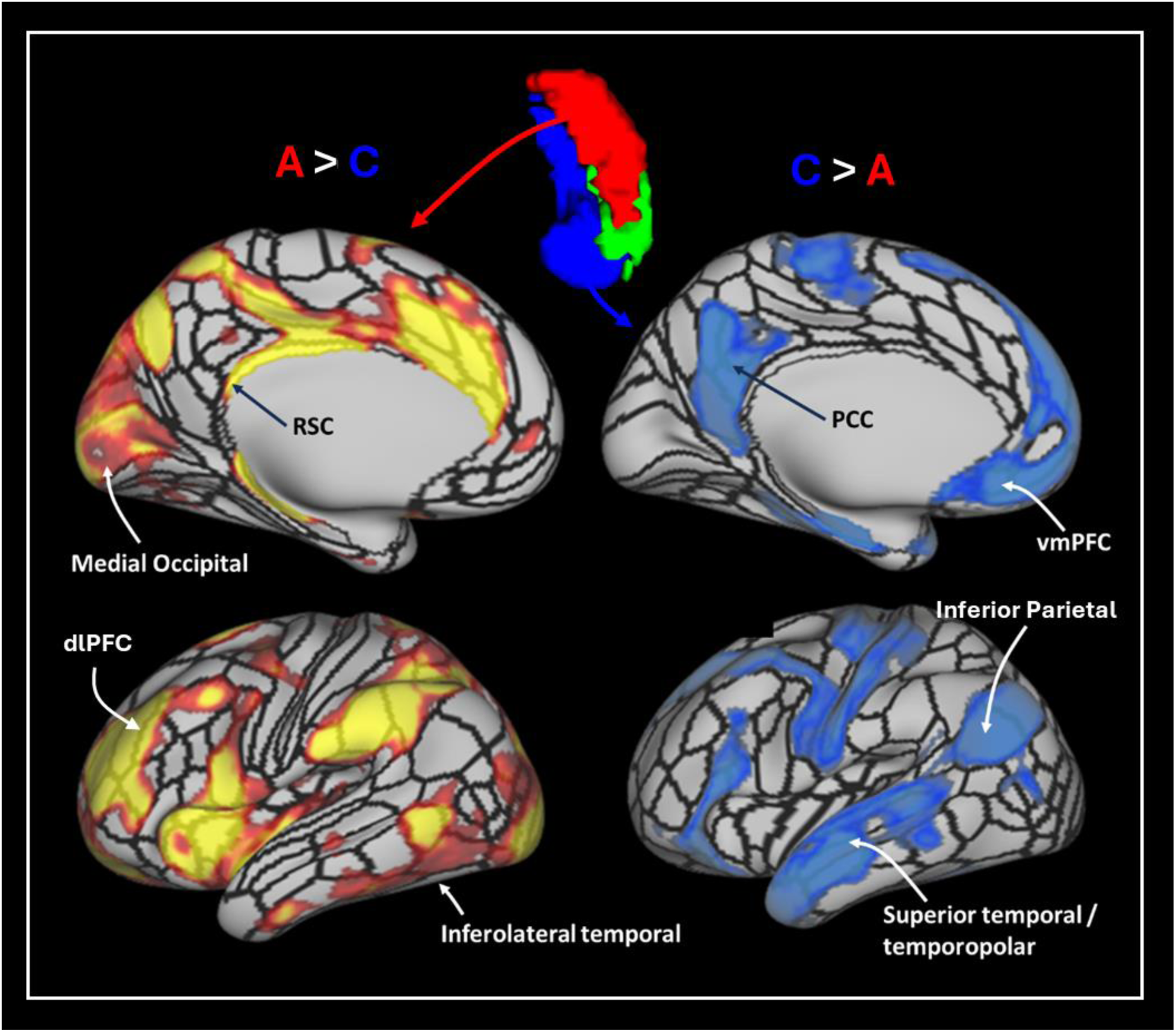
Representative example of differential FC between TW-dFC derived hippocampal clusters and the cortical mantle. Compared with Cluster C, Cluster A displayed preferential FC with RSC, medial occipital visual areas, inferolateral temporal cortex and dlPFC. In contrast, compared with Cluster A, Cluster C displayed preferential FC with PCC, vmPFC, inferior lateral parietal and superior temporal/temporopolar areas.

Similar dissociations were observed in FC patterns between our TW-dFC hippocampal clusters and other cortical areas including occipital, temporal and frontal areas. For example, compared to Clusters B and C, Cluster A exhibited preferential FC with medial occipital and inferolateral temporal areas (see Figure 5; top and bottom left and Supplementary Figures 3 and 4). In contrast, compared to cluster A, Clusters B and C displayed preferential FC with specific regions of the inferior parietal cortex, superior temporal, temporopolar cortices and vmPFC (see Figure 5; bottom right and Supplementary Figures 5 and 7). The interested reader can view the results of all contrasts in Supplementary Figures 3-8 where all statistically significant areas are labelled as defined by the HCPMMP. We now more specifically discuss interactions between our TW-dFC hippocampal clusters and the surrounding medial temporal lobe (MTL) cortices.

FC patterns between TW-dFC derived hippocampal clusters and adjacent MTL cortices revealed equally striking specificity. Compared to Clusters B and C, Cluster A displayed selective FC with a circumscribed region of the lateral entorhinal/medial perirhinal cortex, whereas Clusters B and C showed preferential FC with distinct portions of the medial entorhinal and posterior parahippocampal cortices (Figure 6). Furthermore, direct comparisons between Clusters B and C revealed further dissociation. Cluster B showed stronger FC with a focal region of the lateral entorhinal/medial perirhinal cortex, while Cluster C displayed stronger FC with posterior parahippocampal areas (Figure 6).

**Figure 6.**
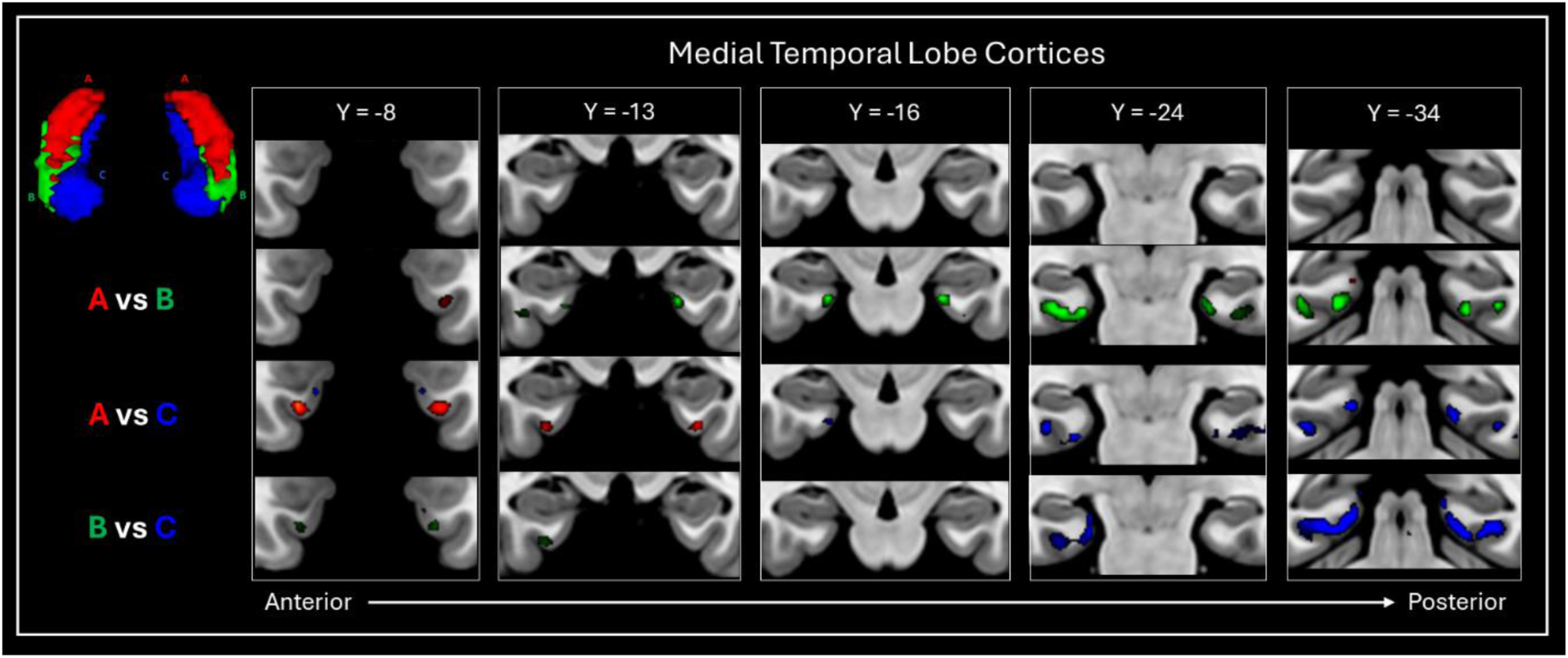
FC patterns between TW-dFC derived hippocampal clusters and surrounding medial temporal (MTL) cortices. Each TW-dFC derived hippocampal cluster displayed specificity in its FC with MTL cortical structures. Compared to Clusters B and C, Cluster A displayed preferentially FC with a distinct region of the lateral entorhinal / medial perirhinal cortex (red). Compared to Cluster A, Clusters B and C showed preferential FC with distinct regions of the medial entorhinal cortex and posterior parahippocampal areas (green and blue). Compared to Cluster C, Cluster B displayed a preferential association with a specific region of the lateral entorhinal / medial perirhinal cortex (green), while Cluster C demonstrated stronger connectivity with posterior parahippocampal areas (blue).

These findings suggest that each TW-dFC derived hippocampal cluster is preferentially coupled with distinct subregions within the MTL cortex, even in the absence of task demands. The spatial precision of these connectivity patterns is notable and, although speculative from this data, may reflect the intrinsic functional architecture that underpins content-specific computations within the MTL, such as those supporting object, spatial or scene processing. This work contributes a novel approach to refine our understanding of intrinsic hippocampal-MTL cortical organisation and provides a foundation for future task-based investigations to more specifically target the functional role of hippocampal-MTL circuits in cognition, including object and spatial memory (Dalton et al., 2018; Grande, Sauvage, Becke, Duzel, & Berron, 2022), navigation (Crivelli-Decker et al., 2023; Sherrill et al., 2013) and related cognitive functions. By integrating anatomically grounded parcellation with dynamic FC analysis, our approach offers a powerful new framework for resolving the fine-scale organisation of hippocampal-MTL cortical interactions. Our methods complement other emerging precision imaging techniques for investigating MTL function (Reznik, Trampel, Weiskopf, Witter, & Doeller, 2023) while offering enhanced anatomical specificity beyond that afforded by conventional functional imaging approaches.

As a final note, distinct FC patterns were also evident between TW-dFC derived hippocampal clusters and subcortical/cerebellar regions. For example, compared to Clusters B and C, Cluster A showed preferential FC with the thalamus, putamen, pallidum, and caudate nucleus (Figure 7). In contrast, compared to Cluster A, Clusters B and C exhibited preferential FC more specifically with the amygdala. Although Clusters B and C each displayed FC with the amygdala compared with Cluster A, Cluster C displayed preferential FC with the amygdala when compared with Cluster B suggesting that Cluster C may have stronger functional links with the amygdala compared with other Clusters. This makes sense considering Cluster C encompassed the uncal region of the hippocampus, which sits adjacent to the hippocampo-amygdalar transition area (HATA). In contrast, compared to Cluster C, Cluster B demonstrated preferential connectivity with the caudate and, to a lesser degree, the putamen and thalamus. Each cluster also displayed preferential connectivity with specific lobes of the cerebellum (Figure 7). These findings offer a more granular understanding of hippocampal-subcortical/cerebellar networks and provide a new approach to investigate how disruptions in these circuits may contribute to the pathophysiology of neuropsychiatric and neurodegenerative disorders.

**Figure 7.**
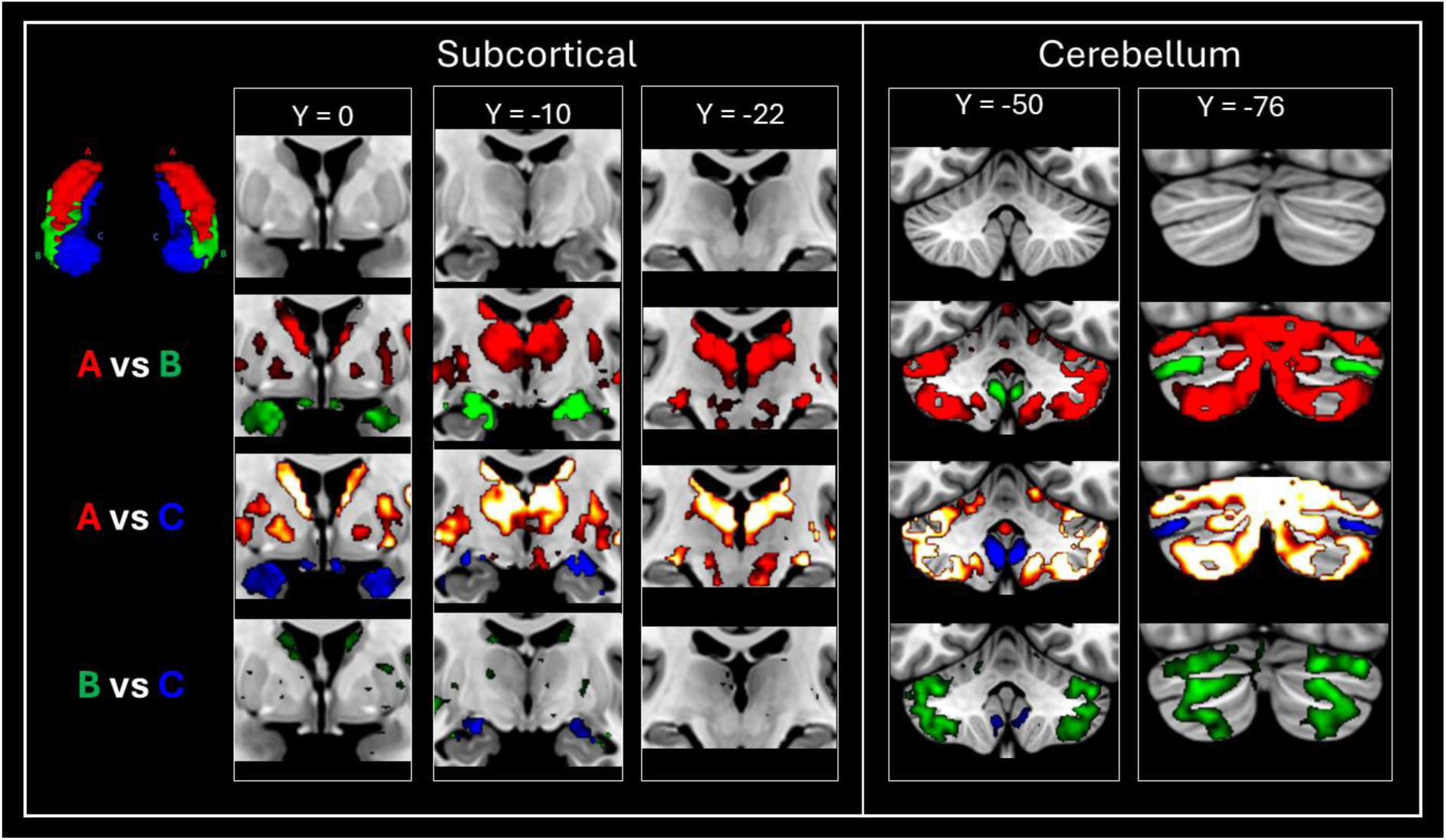
FC patterns between TW-dFC derived hippocampal clusters, subcortical and cerebellar regions. Each TW-dFC derived hippocampal cluster displayed specificity in its FC with subcortical and cerebellar areas. Compared to Clusters B and C, Cluster A showed preferential FC with the thalamus, putamen, pallidum, and caudate nucleus. Compared to Cluster A, Clusters B and C exhibited preferential FC more specifically with the amygdala. Each cluster also displayed preferential connectivity with specific lobes of the cerebellum.

Collectively, the distinct patterns of functional connectivity revealed by our TW-dFC derived hippocampal parcellation highlight its power to delineate how specific hippocampal subdivisions preferentially interact with discrete cortical, subcortical, and cerebellar regions. Extending prior work on anterior-posterior functional differentiation within the hippocampus, this study introduces a novel, anatomically constrained approach to functional parcellation of the hippocampus, grounded in its direct white matter connections with the broader brain. While the present implementation focused on resting-state fMRI, this framework is inherently adaptable to task-based paradigms (Calamante et al., 2017), offering a systematic means to track how hippocampal-cortical interactions may dynamically reconfigure to support diverse cognitive operations (Aly & Turk-Browne, 2018). Our TW-dFC approach lays foundational groundwork for advancing functional models of the hippocampus and its integration within large-scale networks. By anchoring functional connectivity analyses to direct anatomical pathways, this method enables more precise interrogation of the neurobiological substrates underpinning hippocampal interactions with distributed brain regions.

This anatomically informed framework opens new avenues for addressing open questions in the field, such as clarifying how distinct portions of the hippocampus interact with specific subregions within the posterior cingulate cortex during executive, memory, and spatial processing (Alexander, Place, Starrett, Chrastil, & Nitz, 2023; Foster et al., 2023), how they coordinate with inferior parietal and temporal cortices to support different elements of semantic cognition (Humphreys, Lambon Ralph, & Simons, 2021; Kuhnke et al., 2023; Thakral, Madore, & Schacter, 2017) or, more broadly, how anatomically linked (sub)cortico-hippocampal interactions may support memory consolidation (Nadel, Samsonovich, Ryan, & Moscovitch, 2000). In doing so, it offers a powerful platform to refine existing theoretical models of (sub)cortico-hippocampal interactions, including scene construction theory (Dalton & Maguire, 2017; Dalton et al., 2018; Maguire & Mullally, 2013) and the PMAT framework (Reagh & Ranganath, 2023; Ritchey, Libby, & Ranganath, 2015) by providing greater specificity in their predictions regarding how distinct hippocampal subcomponents dynamically interact with distributed (sub)cortical systems. Overall, our findings contribute to ongoing efforts to unravel the complex relationship between hippocampal structure and function, with implications for understanding its role in cognition and its disruption during healthy ageing and in neurological and psychiatric disease.

## Supporting information

Dalton_et_al_Hippocampus_TW-dFC_2025_Supplementary_Video_1

## Acknowledgements

Data were provided by the Human Connectome Project, WU-Minn Consortium (principal investigators: David Van Essen and Kamil Ugurbil; 1U54MH091657) funded by the 16 NIH Institutes and Centers that support the NIH Blueprint for Neuroscience Research; and by the McDonnell Center for Systems Neuroscience at Washington University, St. Louis, MO.

We are grateful for the support of the Australian Research Council (grant numbers DP240102161 and DP210102378). The authors acknowledge the technical assistance provided by the Sydney Informatics Hub and Sydney Imaging, two Core Research Facilities of the University of Sydney, Australia and the facilities and scientific and technical assistance of the National Imaging Facility, a National Collaborative Research Infrastructure Strategy (NCRIS) capability, at Sydney Imaging, the University of Sydney.

---

**Supplementary Table 1.** Results of the Study 2 rsfMRI comparisons. Results are reported at P<0.05 FDR corrected.

|  |  |  | MNI Peak Co-ordinate |  |  |  |
| --- | --- | --- | --- | --- | --- | --- |
| Contrast | Location of Peak Voxel | Hemi-sphere | x | y | z | Number of Voxels |
| Cluster A > Cluster B | Cerebellum | L | -30 | -70 | -56 | 20641 |
|  | Dorsal anterior cingulate gyrus | L | -6 | 20 | 40 | 5101 |
|  | Lateral fronto-orbital gyrus | L | -22 | 50 | -14 | 505 |
|  | Inferior occipital gyrus | L | -50 | -62 | -4 | 342 |
|  | Posterior middle temporal gyrus | R | 58 | -50 | -8 | 318 |
|  | Superior frontal gyrus (posterior segment) | L | -14 | 2 | 68 | 293 |
|  | Middle frontal gyrus (dorsal prefrontal cortex) | R | 24 | 52 | 24 | 173 |
|  | Superior frontal gyrus (posterior segment) | R | 6 | 30 | 44 | 102 |
|  | Fusiform gyrus | L | -32 | -34 | -10 | 26 |
|  | Dorsal anterior cingulate gyrus | L | -2 | 22 | 26 | 24 |
|  | Postcentral gyrus | L | -20 | -32 | 62 | 20 |
| Cluster B > Cluster A |  |  |  |  |  |  |
|  | Precentral gyrus | L | -6 | -24 | 62 | 2514 |
|  | Fusiform gyrus | R | 32 | -32 | -12 | 1976 |
|  | Amygdala | R | -20 | -6 | -26 | 1940 |
|  | Postcentral gyrus | R | 6 | -26 | 58 | 1725 |
|  | Subgenual anterior cingulate gyrus | R | 4 | 12 | -10 | 652 |
|  | Cerebellum | R | 34 | -76 | -34 | 556 |
|  | Cerebellum | L | -14 | -88 | -38 | 387 |
|  | Inferior frontal gyrus pars opercularis | L | -52 | 20 | 18 | 344 |
|  | Cerebellum | R | 6 | -52 | -42 | 294 |
|  | Inferior frontal gyrus pars orbitalis | R | 48 | 36 | -12 | 255 |
|  | Precuneus | R | 6 | -64 | 44 | 139 |
|  | Precuneus | L | -10 | -54 | 34 | 123 |
|  | Posterior cingulate gyrus | R | 16 | -54 | 18 | 97 |
|  | Middle occipital gyrus | L | -36 | -78 | 38 | 85 |
|  | Posterior cingulate gyrus | L | -6 | -54 | 10 | 84 |
|  | Angular gyrus | R | 52 | -50 | 24 | 61 |
|  | Supramarginal gyrus | R | 48 | -18 | 18 | 61 |
|  | Precentral gyrus | R | 40 | -6 | 16 | 40 |
|  | Fusiform gyrus | L | -42 | -48 | -18 | 38 |
|  | Supramarginal gyrus | L | -42 | -26 | 20 | 37 |
|  | Middle occipital gyrus | R | 48 | -70 | 8 | 29 |
|  | Posterior middle temporal gyrus | L | -50 | -54 | 20 | 25 |
|  | Middle occipital gyrus | L | -46 | -76 | 12 | 25 |
| Cluster A > Cluster C | Superior frontal gyrus (posterior segment) | R | 8 | 10 | 48 | 5594 |
|  | Precentral gyrus | L | -28 | -6 | 50 | 140 |
|  | Superior frontal gyrus (posterior segment) | L | -18 | 14 | 62 | 130 |
|  | Anterior cingulate gyrus | R | 6 | 40 | 6 | 105 |
|  | Inferior temporal gyrus | L | -52 | -10 | -32 | 102 |
|  | Inferior temporal gyrus | R | 48 | -2 | -34 | 96 |
|  | Entorhinal/fusiform cortex | R | 24 | -4 | -34 | 53 |
|  | Entorhinal/fusiform cortex | L | -24 | -4 | -36 | 41 |
|  | Postcentral gyrus | L | -20 | -32 | 62 | 29 |
|  | Superior frontal gyrus (prefrontal cortex) | L | -10 | 52 | 2 | 25 |
|  | Rostral anterior cingulate gyrus | L | -4 | 42 | 4 | 22 |
|  | Posterior middle temporal gyrus | L | -52 | -28 | -6 | 20 |
| Cluster C > Cluster A | Precentral gyrus | R | 36 | -22 | 54 | 1518 |
|  | Postcentral gyrus | L | -62 | -10 | 30 | 1063 |
|  | Superior frontal gyrus (prefrontal cortex) | R | 6 | 50 | 22 | 558 |
|  | Postcentral gyrus | L | -10 | -28 | 72 | 388 |
|  | Superior frontal gyrus (prefrontal cortex) | L | -6 | 60 | 18 | 364 |
|  | Cerebellum | R | 6 | -50 | -46 | 348 |
|  | Gyrus rectus | L | -4 | 22 | -12 | 252 |
|  | Cerebellum | R | 40 | -72 | -36 | 167 |
|  | Cerebellum | R | 14 | -86 | -38 | 166 |
|  | Cerebellum | L | -12 | -86 | -40 | 165 |
|  | Inferior frontal gyrus pars triangularis | L | -46 | 24 | 20 | 132 |
|  | Precuneus | R | 6 | -64 | 46 | 122 |
|  | Cerebellum | L | -38 | -72 | -36 | 122 |
|  | Precuneus | L | -8 | -54 | 40 | 103 |
|  | Inferior frontal gyrus pars orbitalis | L | -40 | 38 | -14 | 100 |
|  | Precuneus | L | -16 | -62 | 22 | 56 |
|  | Angular gyrus | R | 52 | -54 | 18 | 54 |
|  | Posterior middle temporal gyrus | L | -60 | -44 | -8 | 53 |
|  | Middle occipital gyrus | R | 38 | -76 | 36 | 52 |
|  | Supramarginal gyrus | R | 42 | -24 | 20 | 48 |
|  | Middle frontal gyrus (posterior segment) | R | 48 | 28 | 18 | 44 |
|  | Inferior frontal gyrus pars orbitalis | R | 38 | 40 | -12 | 41 |
|  | Superior frontal gyrus (posterior segment) | L | -4 | 4 | 66 | 41 |
|  | Inferior frontal gyrus pars orbitalis | R | 42 | 30 | -14 | 30 |
|  | Middle occipital gyrus | L | -46 | -76 | 12 | 29 |
|  | Supramarginal gyrus | L | -44 | -24 | 20 | 28 |
|  | Precentral gyrus | R | 40 | -6 | 16 | 28 |
|  | Subgenual anterior cingulate gyrus | R | 2 | 22 | -8 | 28 |
|  | Posterior middle temporal gyrus | R | 62 | -40 | -8 | 27 |
|  | Superior occipital gyrus | R | 22 | -60 | 22 | 22 |
| Cluster B > Cluster C | Cerebellum | L | -42 | -56 | -44 | 6945 |
|  | Posterior cingulate gyrus | R | 4 | -26 | 40 | 6030 |
|  | Anterior insular | L | -38 | 12 | -10 | 1687 |
|  | Lateral fronto-orbital gyrus | L | -26 | 44 | -12 | 784 |
|  | Supramarginal gyrus | L | -56 | -42 | 40 | 454 |
|  | Angular gyrus | R | 46 | -56 | 46 | 433 |
|  | Middle temporal gyrus | R | 52 | -24 | -8 | 412 |
|  | Posterior middle temporal gyrus | L | -52 | -32 | -4 | 336 |
|  | Anterior middle temporal gyrus | L | -52 | -6 | -30 | 322 |
|  | Inferior temporal gyrus | R | 50 | 4 | -34 | 280 |
|  | Middle occipital gyrus | R | 30 | -92 | 2 | 136 |
|  | Superior occipital gyrus | R | 16 | -64 | 30 | 117 |
|  | Putamen | L | -26 | -2 | 8 | 106 |
|  | Medulla | R | 4 | -40 | -44 | 83 |
|  | Middle occipital gyrus | L | -36 | -88 | -6 | 73 |
|  | Superior parietal gyrus | L | -24 | -48 | 64 | 63 |
|  | Anterior fusiform gyrus | R | 24 | -8 | -36 | 47 |
|  | Cuneus | L | -6 | -82 | 34 | 36 |
|  | Pallidum | R | 16 | -4 | -4 | 35 |
|  | Superior parietal gyrus | R | 28 | -44 | 64 | 33 |
|  | Brain stem, pontine crossing tract | R | 6 | -24 | -28 | 32 |
|  | Cuneus | R | 12 | -80 | 12 | 29 |
|  | Cerebellum | R | 28 | -34 | -37 | 26 |
|  | Cuneus | L | -12 | -74 | 42 | 24 |
| Cluster C > Cluster B | Posterior cingulate gyrus | L | -6 | -56 | 12 | 271 |
|  | Gyrus rectus (vmPFC) | L | -8 | 46 | -10 | 265 |
|  | Posterior cingulate gyrus | R | 6 | -58 | 16 | 264 |
|  | Posterior fusiform gyrus | L | -24 | -32 | -20 | 246 |
|  | Gyrus rectus (vmPFC) | R | 4 | 38 | -18 | 228 |
|  | Posterior fusiform gyrus | R | 28 | -30 | -18 | 227 |
|  | Superior frontal gyrus (posterior segment) | L | -22 | 28 | 40 | 221 |
|  | Superior frontal gyrus (posterior segment) | R | 26 | 20 | 46 | 162 |
|  | Middle occipital gyrus | L | -38 | -80 | 32 | 158 |
|  | Angular gyrus | R | 44 | -70 | 36 | 133 |
|  | Parahippocampal gyrus / amygdala | L | -22 | -4 | -28 | 128 |
|  | Amygdala | R | 18 | -8 | -17 | 124 |
|  | Cerebellum | L | -10 | -86 | -38 | 113 |
|  | Middle temporal gyrus | R | 54 | -6 | -18 | 109 |
|  | Precentral gyrus | R | 6 | -26 | 62 | 91 |
|  | Middle frontal gyrus (posterior segment) | L | -44 | 28 | 18 | 79 |
|  | Cerebellum | R | 7 | -51 | -47 | 79 |
|  | Inferior frontal gyrus pars orbitalis | L | -38 | 34 | -12 | 71 |
|  | Cerebellum | L | -8 | -46 | -42 | 63 |
|  | Middle temporal gyrus | L | -60 | -8 | -18 | 63 |
|  | Lateral fronto-orbital gyrus | R | 34 | 34 | -12 | 60 |
|  | Cerebellum | R | 14 | -86 | -36 | 51 |
|  | Precentral gyrus | L | -10 | -20 | 72 | 37 |
|  | Posterior middle temporal gyrus | L | -58 | -46 | -10 | 35 |
|  | Posterior fusiform gyrus | R | 42 | -28 | -22 | 33 |
|  | Middle frontal gyrus (posterior segment) | R | 44 | 32 | 18 | 30 |
|  | Postcentral gyrus | L | -4 | -34 | 60 | 26 |
|  | Pole of middle temporal gyrus | R | 42 | 16 | -34 | 24 |
|  | Postcentral gyrus | R | 42 | -14 | 56 | 22 |

## Appendices

**Supplementary Video 1. Representative TW-dFC map from a single participant.** This video provides a compelling visualisation of time-resolved hippocampal connectivity, capturing dynamic fluctuations in how the hippocampus functionally engages with the broader brain over time. Functional coupling is projected onto the white matter pathways that mediate these interactions, illustrating how specific anatomical tracts mediate shifting patterns of communication between the hippocampus and distributed cortical and subcortical regions. Notably, the hippocampus exhibits time-varying interactions with occipital, parietal, temporal, frontal, and thalamic areas throughout the resting-state scan. This dynamic mapping underscores the temporally evolving nature of hippocampal network integration, shaped and constrained by its structural connectivity.

**Supplementary Figure 1.**
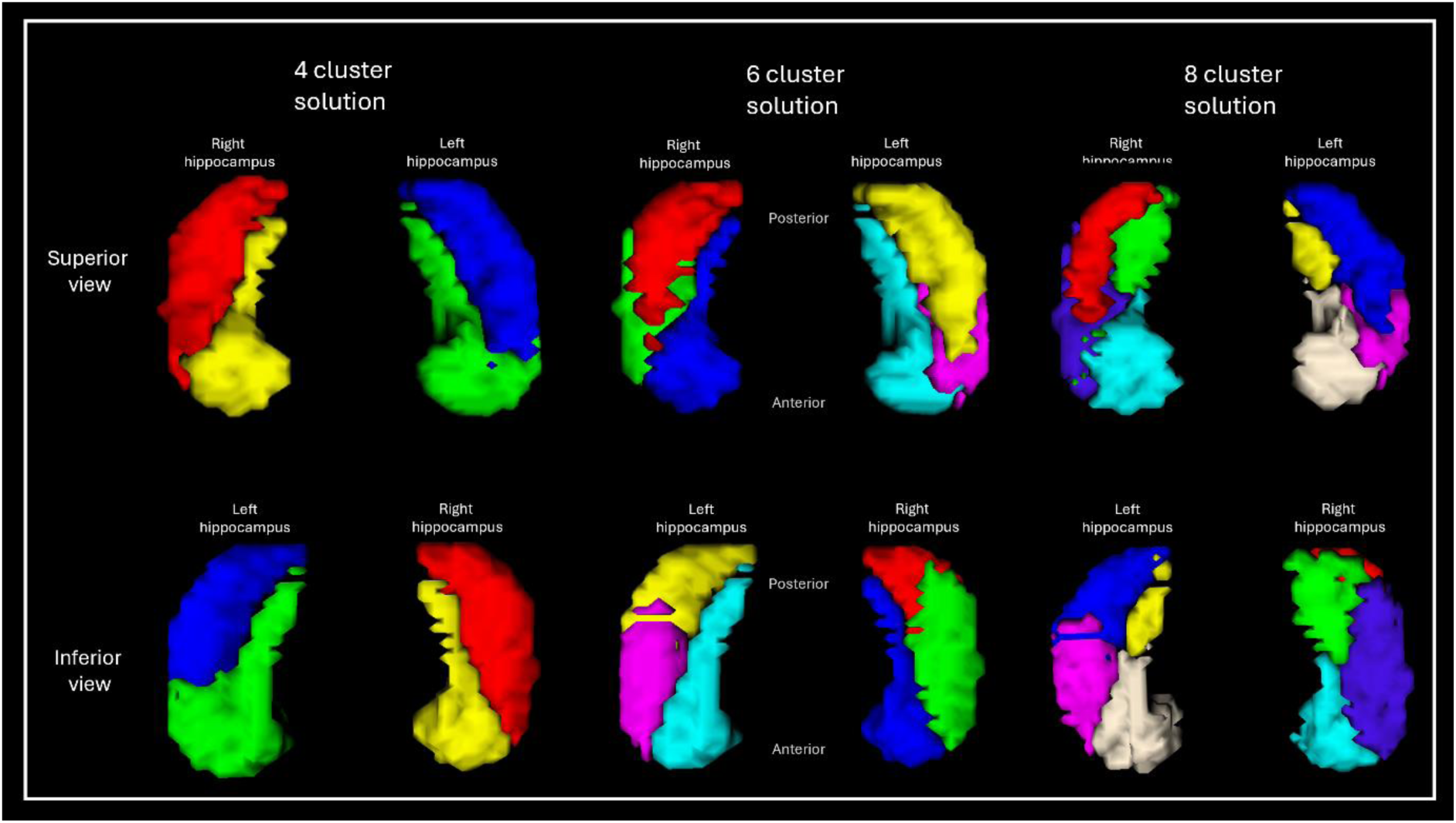
3D renderings of hippocampal clusters identified via k-means clustering of independent component analysis (ICA) on TW-dFC data, shown for 4-, 6-, and 8-cluster solutions. Clusters are distributed along both the anterior-posterior (anterior towards the bottom of the figure, posterior towards the top of the figure) and medial-lateral axes of the hippocampus. 4-cluster solution (left); Rather than a simple anterior-posterior split, this solution revealed two spatially distinct divisions within each hippocampus: an anteromedial division encompassing the uncal and medial portions (yellow and green clusters), and a posterolateral division encompassing the hippocampal tail and lateral portions (red and blue clusters). 6-cluster solution (middle); This parcellation is described in detail in the main text and Figure 2. 8-cluster solution (right); Similar in organisation to the 6-cluster solution but displayed a further subdivision of the medial cluster into anterior (cyan and beige) and posterior (green and yellow) components. Across all solutions, note the striking bilateral symmetry in cluster organisation across the left and right hippocampus, despite no spatial constraints having been imposed during clustering.

**Supplementary Figure 2.**
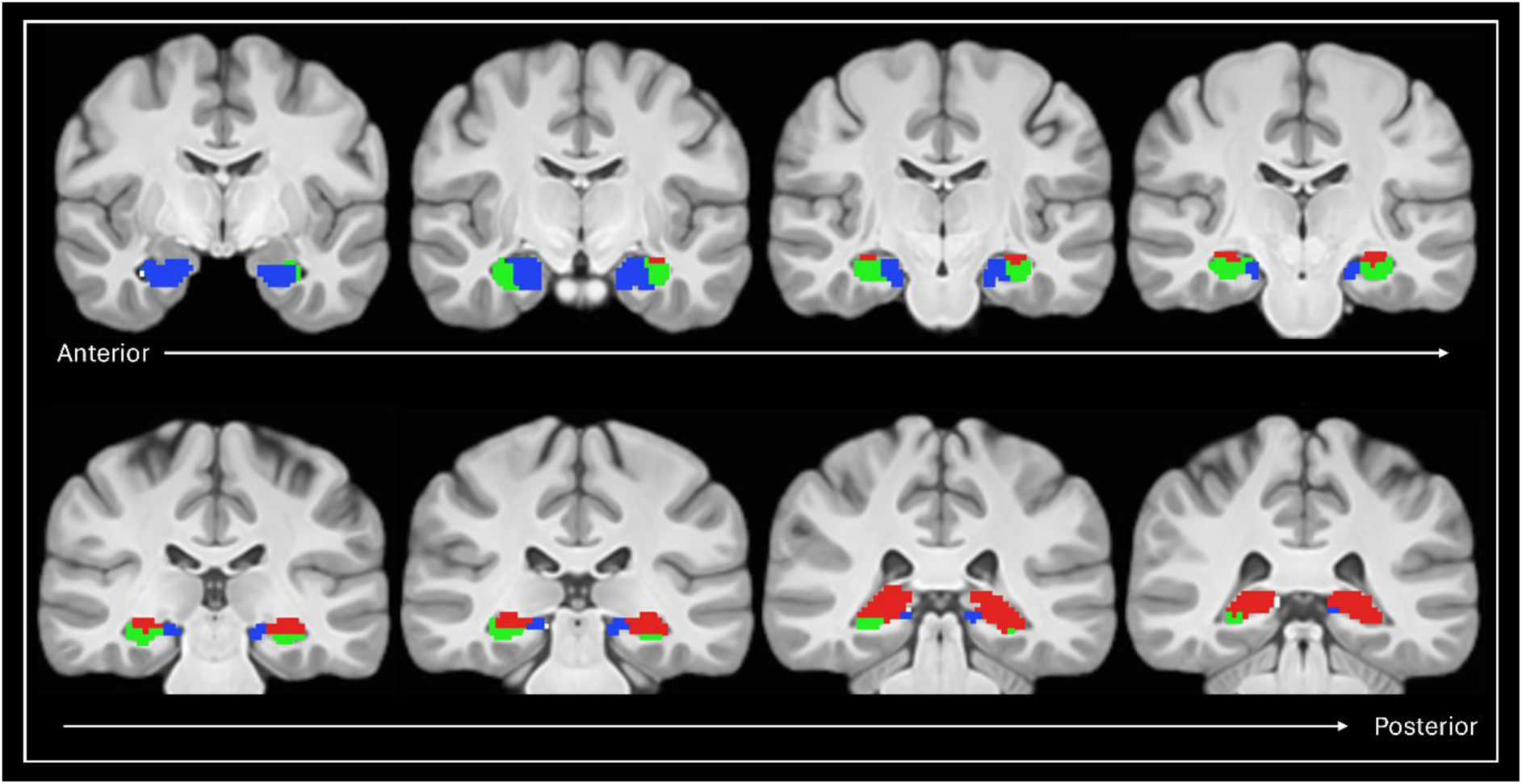
Anterior-posterior distribution of TW-dFC-derived clusters. Clusters identified using the 6-cluster TW-dFC solution are overlaid on coronal slices of a T1-weighted image progressing from the anterior (top left) to posterior (bottom right) extent of the hippocampus. Within the hippocampal body, individual clusters show broad spatial correspondence with cytoarchitecturally defined subfields (described in text and visually presented in Figure 2B). In contrast, broader functional clusters were observed in the hippocampal head and tail, encompassing multiple subfields and suggesting coarser functional organisation in these regions.

**Supplementary Figure 3.**
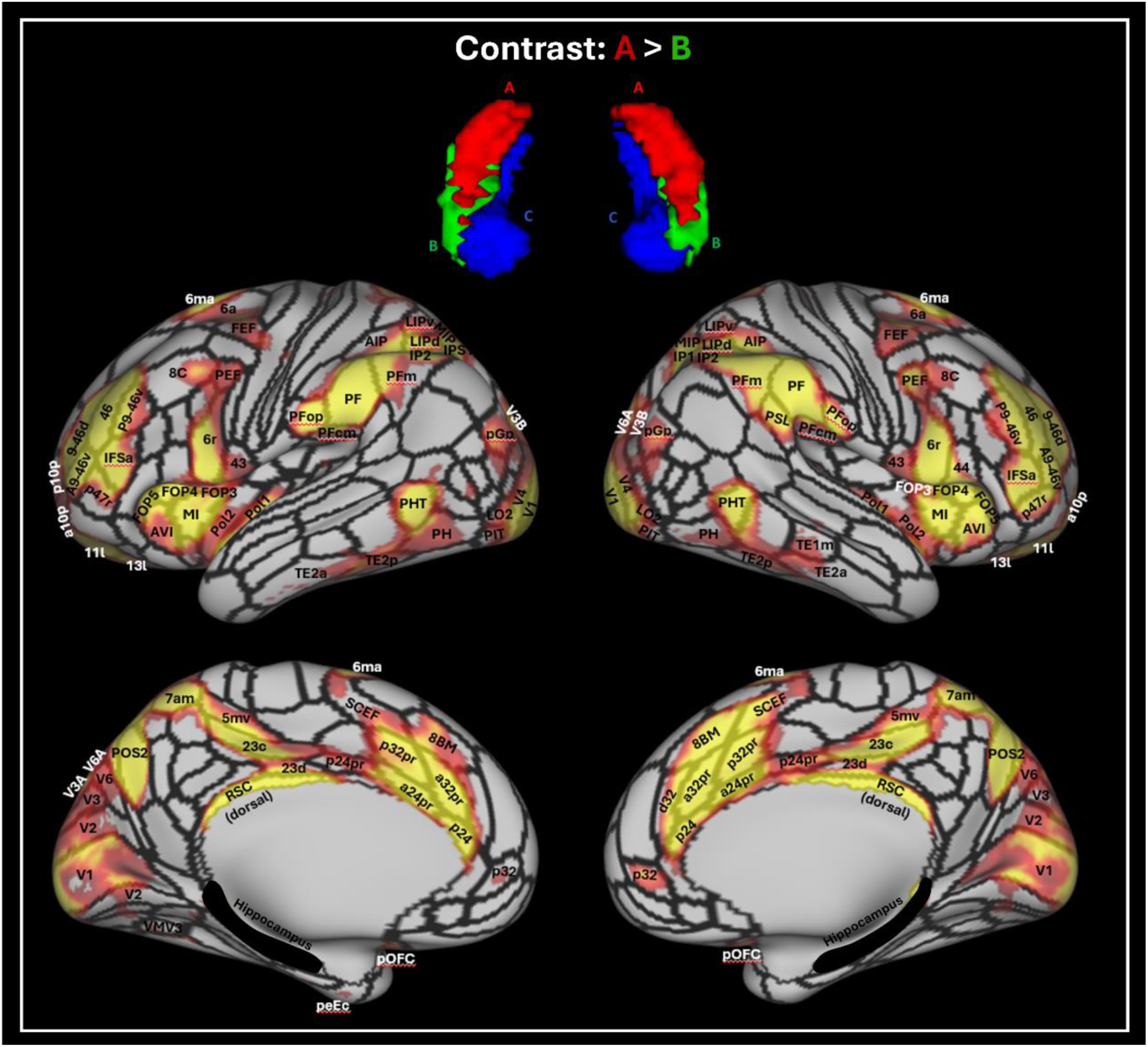
Results of the seed-based rsfMRI analysis based on the TW-dFC hippocampal clusters. Results for the contrast of cluster A (hippocampal tail cluster) > cluster B (anterolateral cluster). Results are overlaid on the HCPMMP (hippocampus is highlighted in black) and statistically significant areas are labelled. T-test results are thresholded at p<0.05 FDR corrected.

**Supplementary Figure 4.**
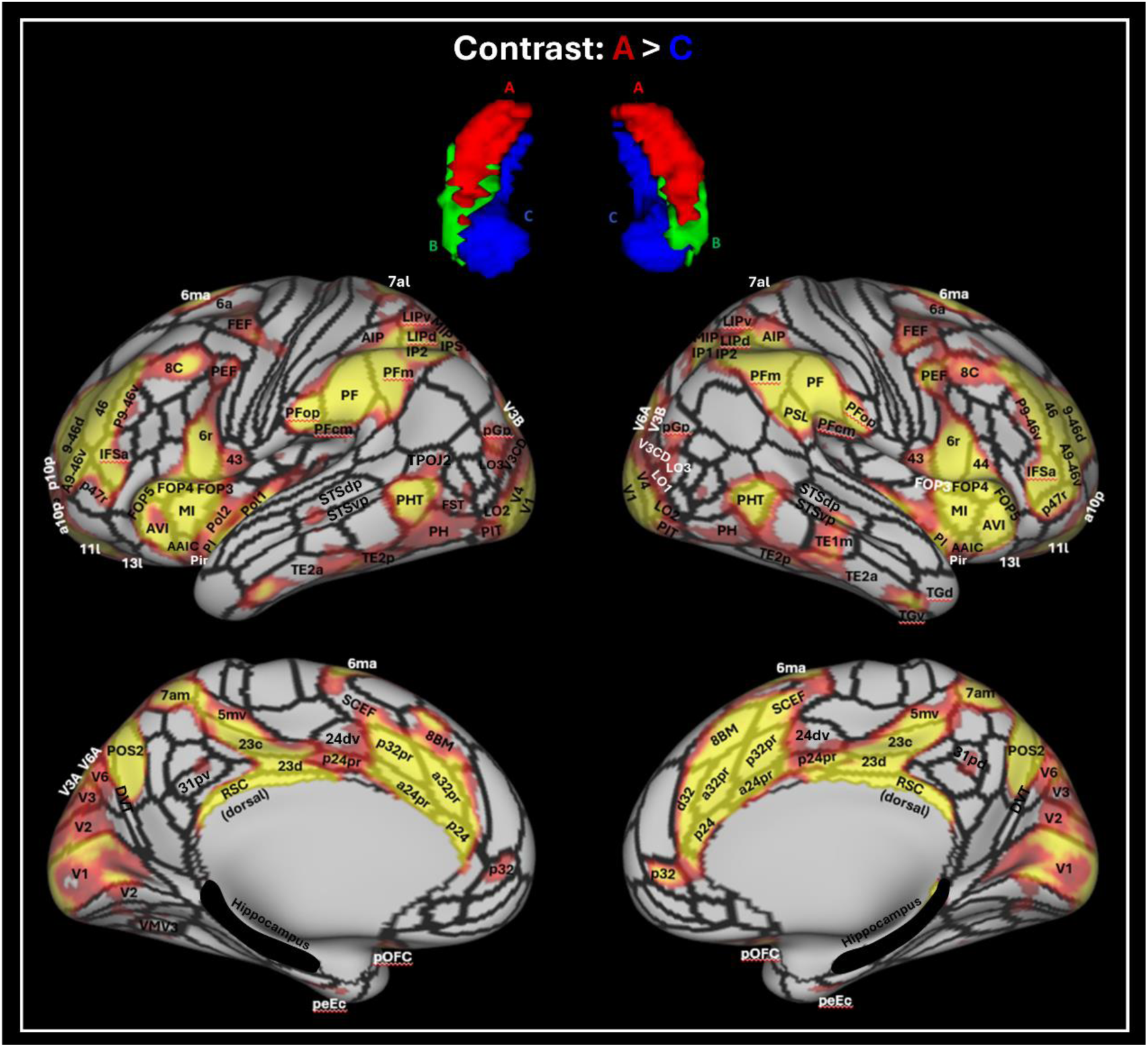
Results of the seed-based rsfMRI analysis based on the TW-dFC hippocampal clusters. Results for the contrast of cluster A (hippocampal tail cluster) > cluster C (medial cluster). Results are overlaid on the HCPMMP (hippocampus is highlighted in black) and statistically significant areas are labelled. T-test results are thresholded at p<0.05 FDR corrected.

**Supplementary Figure 5.**
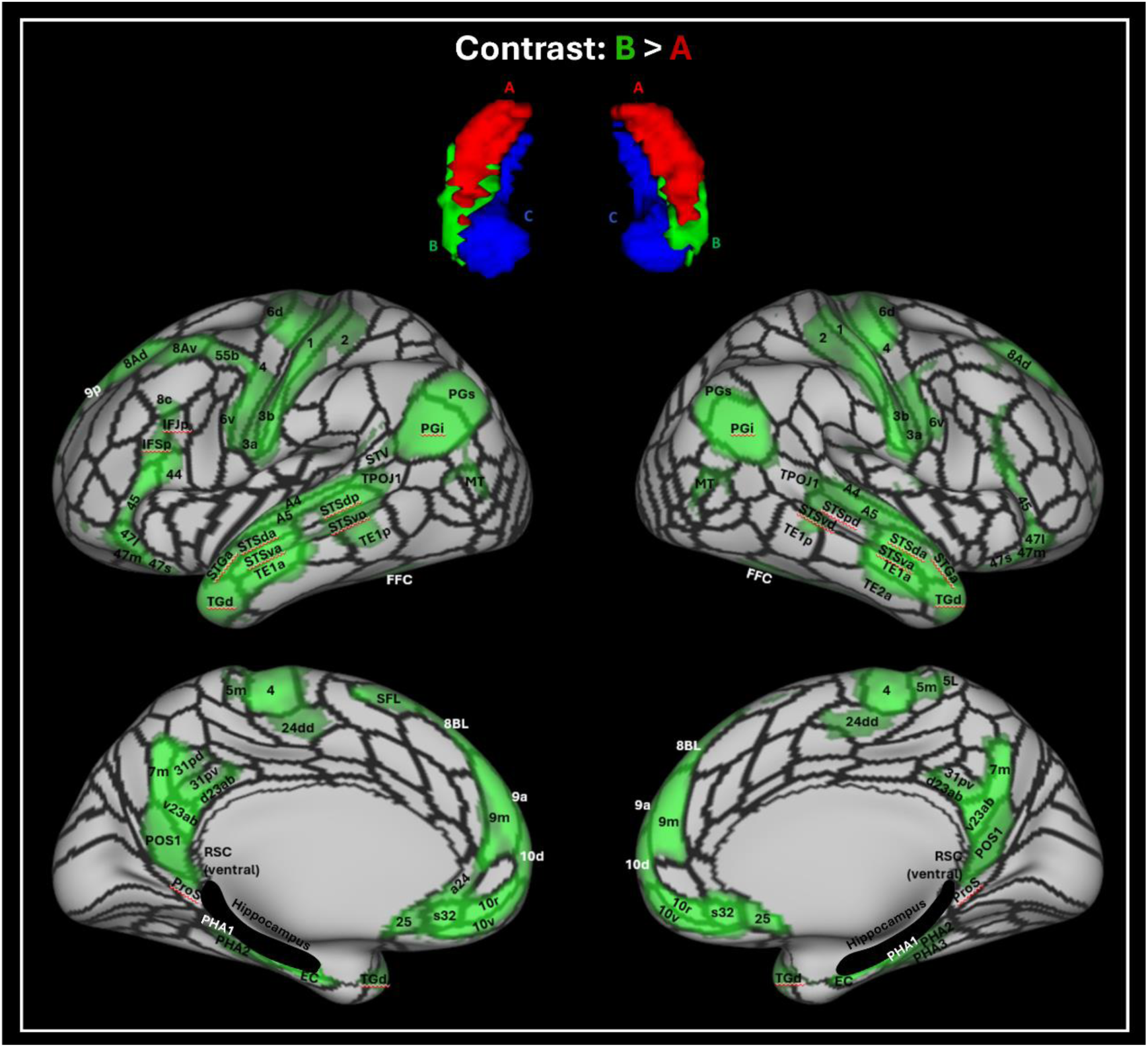
Results of the seed-based rsfMRI analysis based on the TW-dFC hippocampal clusters. Results for the contrast of cluster B (anterolateral cluster) > cluster A (hippocampal tail cluster). Results are overlaid on the HCPMMP (hippocampus is highlighted in black) and statistically significant areas are labelled. T-test results are thresholded at p<0.05 FDR corrected.

**Supplementary Figure 6.**
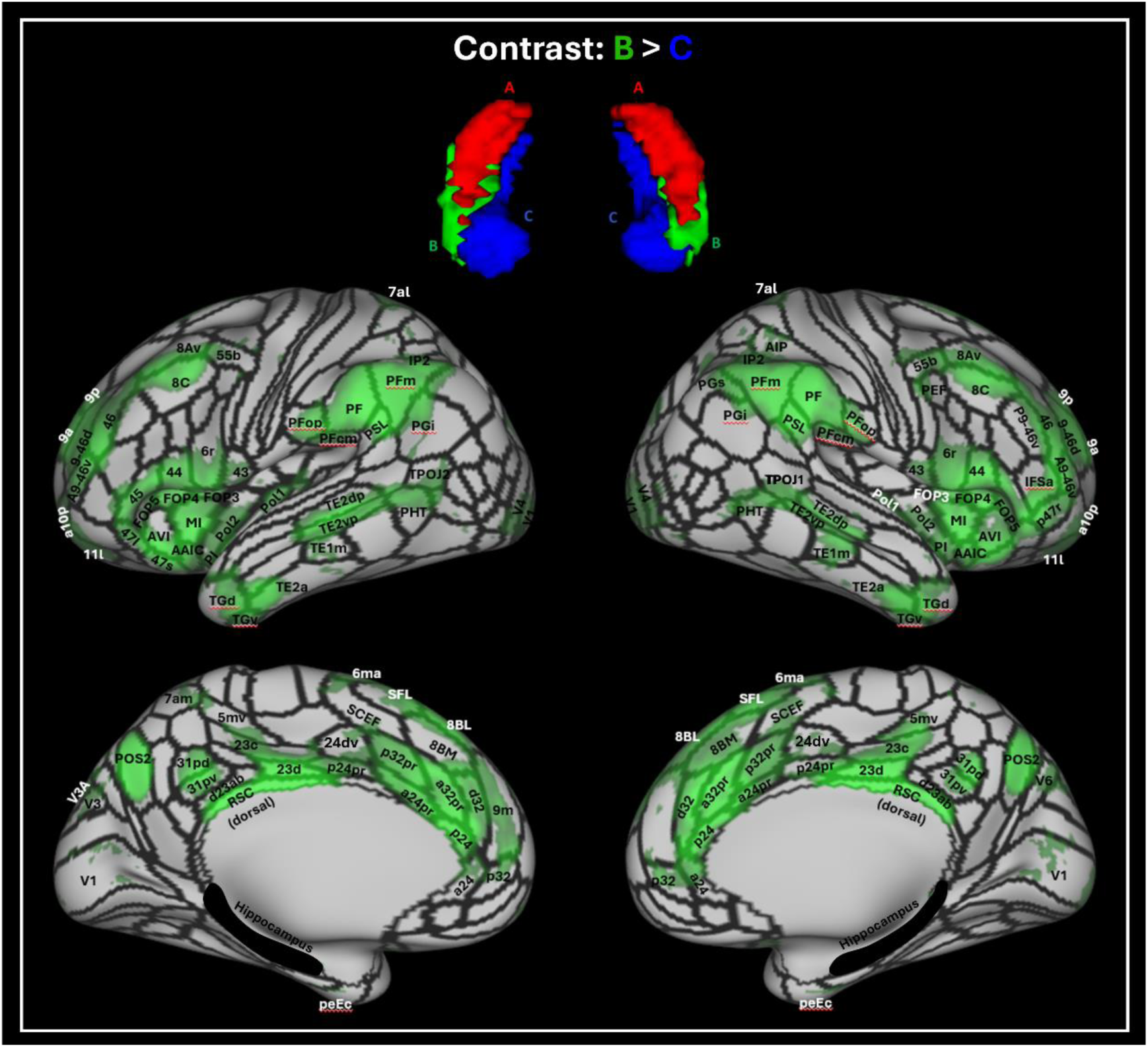
Results of the seed-based rsfMRI analysis based on the TW-dFC hippocampal clusters. Results for the contrast of cluster B (anterolateral cluster) > cluster C (medial cluster). Results are overlaid on the HCPMMP (hippocampus is highlighted in black) and statistically significant areas are labelled. T-test results are thresholded at p<0.05 FDR corrected.

**Supplementary Figure 7.**
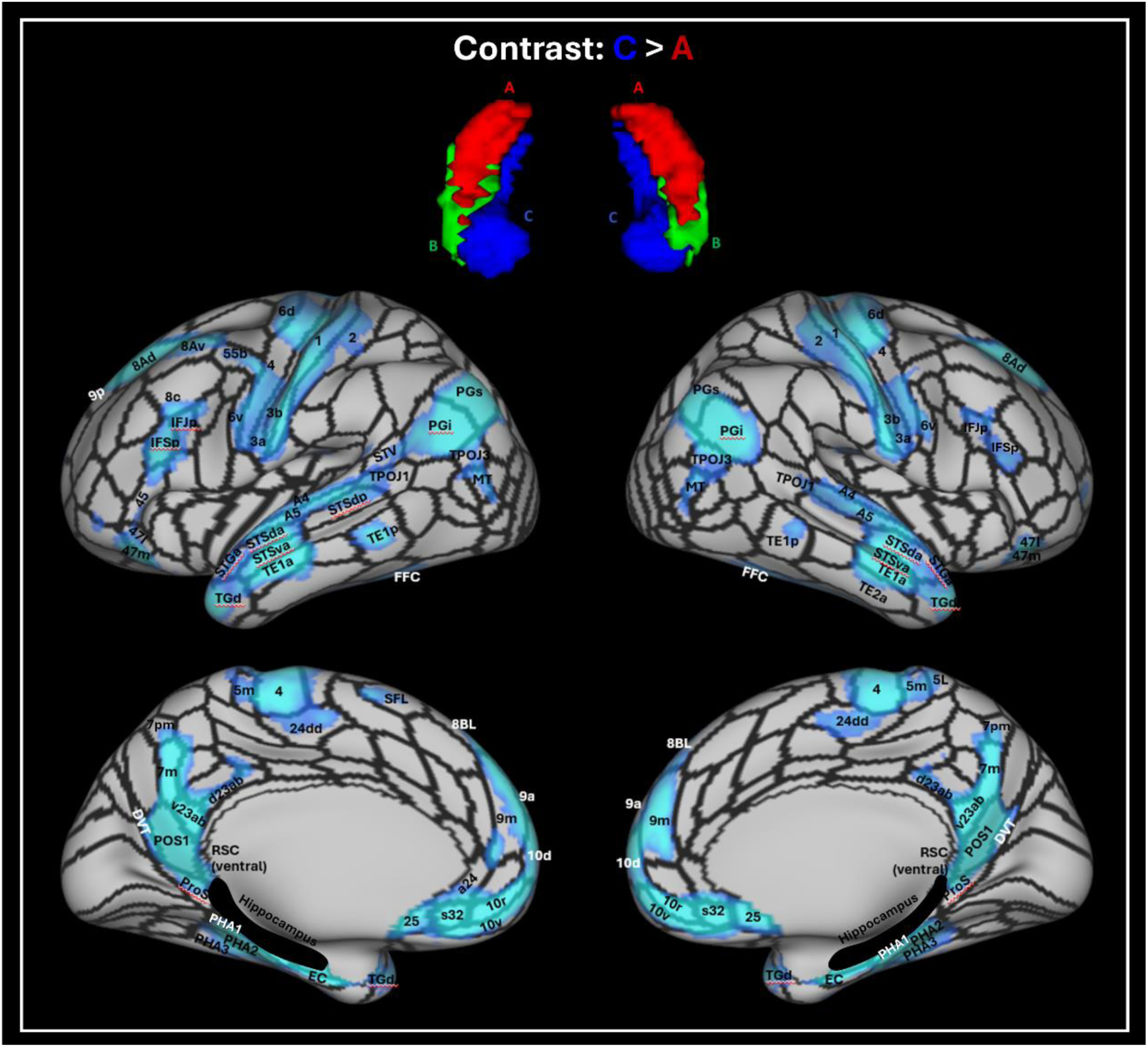
Results of the seed-based rsfMRI analysis based on the TW-dFC hippocampal clusters. Results for the contrast of cluster C (medial cluster) > cluster A (hippocampal tail cluster). Results are overlaid on the HCPMMP (hippocampus is highlighted in black) and statistically significant areas are labelled. T-test results are thresholded at p<0.05 FDR corrected.

**Supplementary Figure 8.**
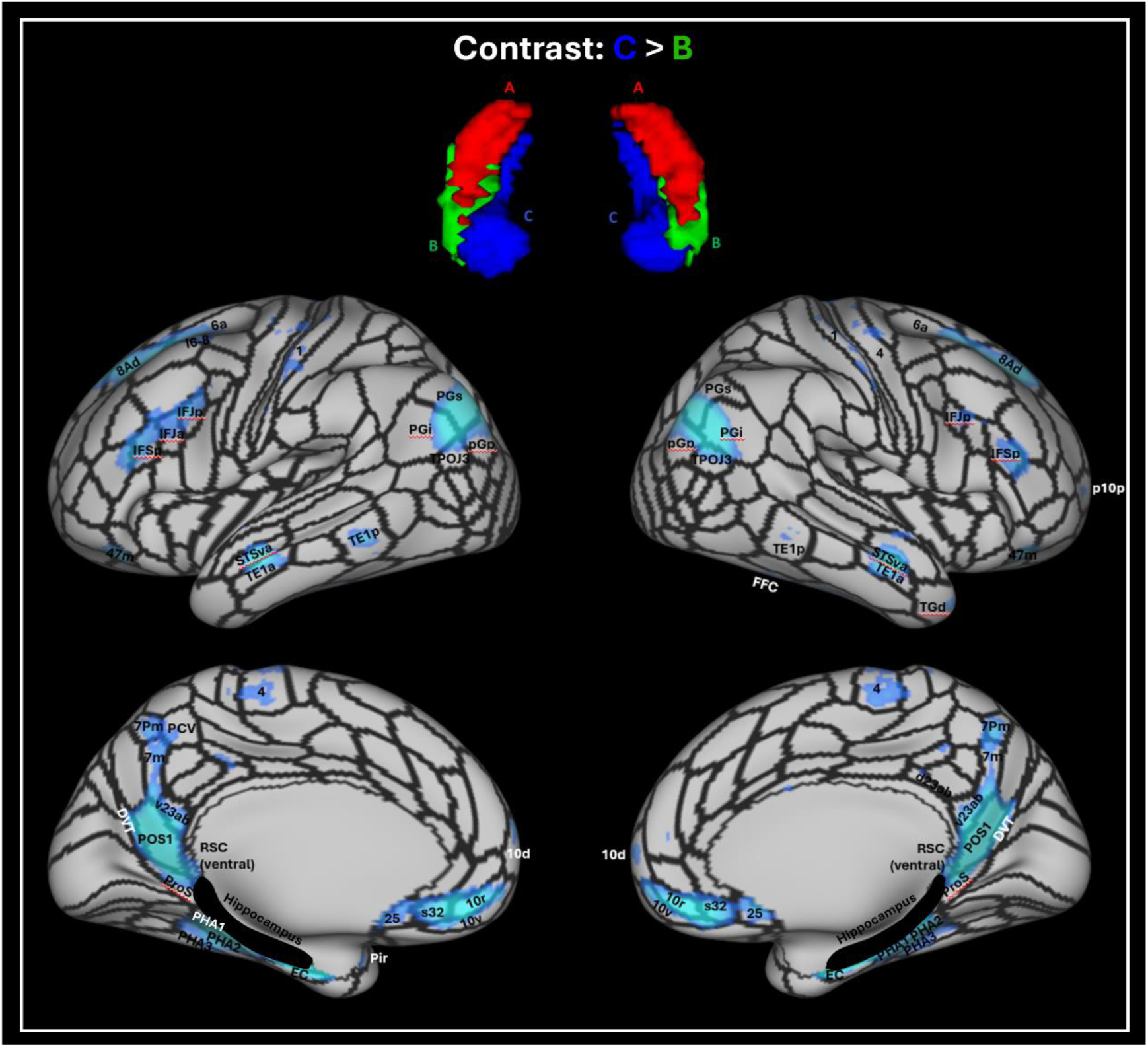
Results of the seed-based rsfMRI analysis based on the TW-dFC hippocampal clusters. Results for the contrast of cluster C (medial cluster) > cluster B (anterolateral cluster). Results are overlaid on the HCPMMP (hippocampus is highlighted in black) and statistically significant areas are labelled. T-test results are thresholded at p<0.05 FDR corrected.

